# Telomere-loop dynamics in chromosome end protection

**DOI:** 10.1101/279877

**Authors:** David Van Ly, Ronnie Ren Jie Low, Sonja Frölich, Tara K. Bartolec, Georgia R. Kafer, Hilda A. Pickett, Katharina Gaus, Anthony J. Cesare

## Abstract

We used super-resolution microscopy to investigate the role of macromolecular telomere structure in chromosome end protection. In murine and human cells with reduced TRF2, we find that ATM-activation at chromosome ends occurs with a structural change from t-loops to linearized chromosome ends through t-loop unfolding. Comparably, we find Aurora B kinase regulates telomere linearity concurrent with ATM activation at telomeres during mitotic arrest. Using a separation of function allele, we find that the TRFH domain of TRF2 regulates t-loop formation while suppressing ATM activity. Notably, we demonstrate that telomere linearity and ATM activation occur separately from telomere fusion via non-homologous end-joining (NHEJ). Further, we show that linear DDR-positive telomeres can remain resistant to fusion, even during an extended G1-arrest when NHEJ is most active. Collectively, these results suggest t-loops act as conformational switches that regulate ATM activation at chromosome ends independent of mechanisms to suppress chromosome end fusion.

## INTRODUCTION

Telomeres are the protective nucleoprotein structures at eukaryotic chromosome termini. Through their position at naturally occurring chromosome ends, telomeres have evolved a specialized relationship with the DNA damage response (DDR) and DNA repair pathways that engage with genomic double strand breaks (Arnoult and Karlseder, 2015). Compromised telomere protection in normal mammalian tissues regulates cellular aging by activating the telomere DDR as a critical and potent tumor suppressor program (Roake and Artandi, 2017). However, illicit DNA repair at telomeres is often deleterious, resulting in end-to-end chromosome fusions and genome instability (Maciejowski et al., 2015), or improper homologous recombination and cellular immortality via alternative lengthening of telomeres (Pickett and Reddel, 2015). It is thus imperative for telomeres to enable DDR signaling at chromosome ends to facilitate their role in tumor suppression, whilst simultaneously preventing illegitimate DNA repair to maintain genome stability. Determining how telomeres manage this coordinated balance of DDR activation and DNA repair inhibition, is critically important to understand telomere-dependent regulation of cellular aging and tumor suppression.

Mammalian telomeric DNA consists of many kilobase pairs (kb) of duplex 5’-T2AG3-3’ repeats that terminate in a 50-400 nucleotide (nt) overhang of the G-rich sequence. DDR and DNA repair activity at chromosome ends is regulated by the telomere-specific “shelterin” protein complex, which binds directly to the telomeric DNA (Lazzerini-Denchi and Sfeir, 2016). Several elegant studies have utilized induced telomere dysfunction through deletion of shelterin components to reveal how each shelterin subunit interacts with DDR and DNA repair pathways (Celli and de Lange, 2005; Denchi and de Lange, 2007; Hockemeyer et al., 2006; Martinez et al., 2009; Sfeir and de Lange, 2012; Sfeir et al., 2009; Takai et al., 2011; Wu et al., 2006). Within shelterin, telomeric repeat binding factor 2 (TRF2) is unique in suppressing both ataxia telangiectasia mutated (ATM) activation at chromosome ends and NHEJ-dependent telomere-telomere fusions (Denchi and de Lange, 2007; Smogorzewska et al., 2002). Strikingly, TRF2 deletion results in ATM activation and the eventual covalent fusion of all the chromosome ends within a cell (Celli and de Lange, 2005).

Two pathways of physiological telomere DDR activation in human cells include: the canonical telomere DDR that is activated during cellular aging through progressive telomere erosion (d’Adda di Fagagna et al., 2003; Harley et al., 1990), and the non-canonical telomere DDR induced during mitotic arrest (Hayashi et al., 2012). Together, the canonical and non-canonical mechanisms of telomere deprotection play a central role in regulating telomere-dependent tumor suppression through replicative senescence and crisis (d’Adda di Fagagna et al., 2003; Hayashi et al., 2015; Herbig et al., 2004).

Physiological telomere deprotection proceeds through an indeterminate “intermediate-state", so-called because the telomere DDR is activated without accompanying telomere fusions (Cesare et al., 2009; Kaul et al., 2012). Observation of aged and cancer cells (Cesare et al., 2009; Kaul et al., 2012), and directed molecular biology experimentation (Benarroch-Popivker et al., 2016; Cesare et al., 2013; Fumagalli et al., 2012; Okamoto et al., 2013), suggest that TRF2 regulates intermediate-state telomeres by independently inhibiting ATM and NHEJ. TRF2 inhibits NHEJ by suppressing RNF168 at γ-H2AX labeled telomeres via its iDDR motif in the hinge domain (Okamoto et al., 2013). ATM inhibition by TRF2 requires its TRFH domain and involves alteration of telomere DNA topology (Benarroch-Popivker et al., 2016; Okamoto et al., 2013). Partial TRF2 depletion induces intermediate-state telomeres, indicating less TRF2 protein occupancy at chromosome ends is required to inhibit NHEJ as compared to ATM (Cesare et al., 2013). Correspondingly, the ATM-dependent telomere DDR induced during mitotic arrest results from partial TRF2 removal from mitotic telomeres, independent of telomere length (Hayashi et al., 2012). DDR-positive telomeres do not fuse during mitotic arrest, consistent with global NHEJ suppression during cell division, and remain fusion resistant when passed into the subsequent G1-phase (Hayashi et al., 2012; Orthwein et al., 2014).

Telomere macromolecular structure may provide an additional measure of regulatory control over DDR activation and DNA repair at chromosome ends. Telomere-loops (t-loops) are lariat structures where the 3’G-rich telomeric DNA overhang is sequestered within a displacement loop in the duplex telomeric DNA of the same chromosome end (Griffith et al., 1999). T-loops were originally identified in mammals (Griffith et al., 1999), and subsequently found in the telomeres of protist, fungal, plant, nematode and avian species (Cesare et al., 2003; Munoz-Jordan et al.,2001; Murti and Prescott, 1999; Nikitina and Woodcock, 2004; Raices et al., 2008). Sequestration of the telomere overhang, and conservation in diverse eukaryotic species, implied protective roles for t-loops at chromosome ends. However, studies of putative t-loop functions were hindered for more than a decade after their discovery, due to the requirement for specialized DNA purification and electron microscopy techniques to visualize these structures.

A major advance in the ability to study telomere macromolecular structure was provided by the de Lange and Zhuang laboratories, who collaborated to develop a technique to visualize t-loops in the elongated telomeres of mouse cells using super-resolution stochastic optical reconstruction microscopy (STORM) (Doksani et al., 2013). Utilizing a panel of shelterin knockout mouse embryonic fibroblasts (MEFs), these experiments revealed that TRF2 is the only shelterin component that contributes to t-loop formation (Doksani et al., 2013), consistent with *in vitro* biochemical experiments implicating TRF2 in establishing t-loops (Griffith et al., 1999; Stansel et al., 2001). In these experiments, telomeres from *TRF2* null cells were examined by STORM 156 hours after gene deletion, when almost all chromosome ends were fused. These findings revealed that t-loops were absent at covalent telomere-telomere fusions, and concluded t-loops simultaneously function to repress ATM and NHEJ activity at chromosome ends (Doksani et al., 2013). A limitation of this study, however, was that telomeres were only observed when chromosome ends were protected from the DDR, at the end point after NHEJ-dependent fusion, or at late time points after TRF2 deletion in ATM-null cells when the telomere DDR was inactivated. Telomeres with an active ATM-dependent DDR were not observed, and the possibility that telomere protection against ATM and NHEJ occurs through independent mechanisms was not considered.

Here we explore the role of telomere macromolecular structure through the continuum of telomere deprotection, including both DDR activation and telomere fusion. In addition to STORM, we describe methods to visualize murine and human t-loops using more rapid methods of super-resolution microscopy. We identify that the ATM-dependent telomere DDR correlates specifically with a telomere structural change from looped to linear telomeres through t-loop unfolding; that TRF2 inhibits ATM activity and regulates t-loop formation via its TRFH domain; and that Aurora B kinase regulates telomere linearization corresponding with activation of the non-canonical mitotic arrest-dependent telomere DDR. We also demonstrate that linear DDR-positive telomeres can remain resistant to chromosome end fusions, even during an extended G1-phase when NHEJ is most active. Cumulatively, these data identify that intermediate-state telomeres are linearized DDR-positive chromosome ends, suggesting that t-loops function as conformational switches to govern ATM-dependent DDR activation at chromosome ends, independently of telomere mechanisms to inhibit NHEJ.

## RESULTS

### Intermediate-state telomeres are conserved in mice

Laboratory mouse telomeres are substantially longer than human telomeres, enabling more efficient t-loop visualization. To determine how telomere structure corresponded with progressive telomere deprotection in MEFs, we had to first determine if intermediate-state telomeres (i.e. DDR-positive/NHEJ-resistant) were conserved in mice. To test this, we developed a temporally controlled experiment following TRF2 deletion in *TRF2^Floxed/Floxed^ Rosa26-CreERT2 pBabeSV40LT* MEFs (hereafter abbreviated *TRF2^F/F^ CreER LgT*, Figure S1A) (Okamoto et al., 2013). Our prediction was that following 4-hydroxytamoxifen (4-OHT)-induced *TRF2* deletion by Cre-ERT2, the gradual turnover of remaining TRF2 protein would facilitate transition of telomeres from a protected state, through the intermediate-state, before sufficient TRF2 was lost to induce telomere fusions. Following 4-OHT treatment, we quantified DDR-positive telomeres, termed “telomere deprotection induced foci” or “TIF” (Takai et al., 2003), in mitotic and interphase cells and measured telomere fusions in cytogenetic chromosome spreads. Metaphase-TIF emerged with 18-24 hours of 4-OHT, and peaked at 48 hours, whereas low numbers of telomere fusions emerged by 48 hours and accumulated over the 120-hour experiment (Figures 1A, B, and S1B-D). Similar TIF kinetics were observed in interphase cells (Figures 1A-D). Notably, DDR foci were resolved at telomere fusions, corresponding with a reduction in interphase-TIF at later time points in the experiment (Figures 1A-D). Previous reports demonstrated that *TRF2* deletion in MEFs induces an ATM-specific telomere DDR (Denchi and de Lange, 2007), indicating the ATM response at mouse chromosome ends is suppressed after NHEJ-dependent telomere fusion.

**Figure 1.**
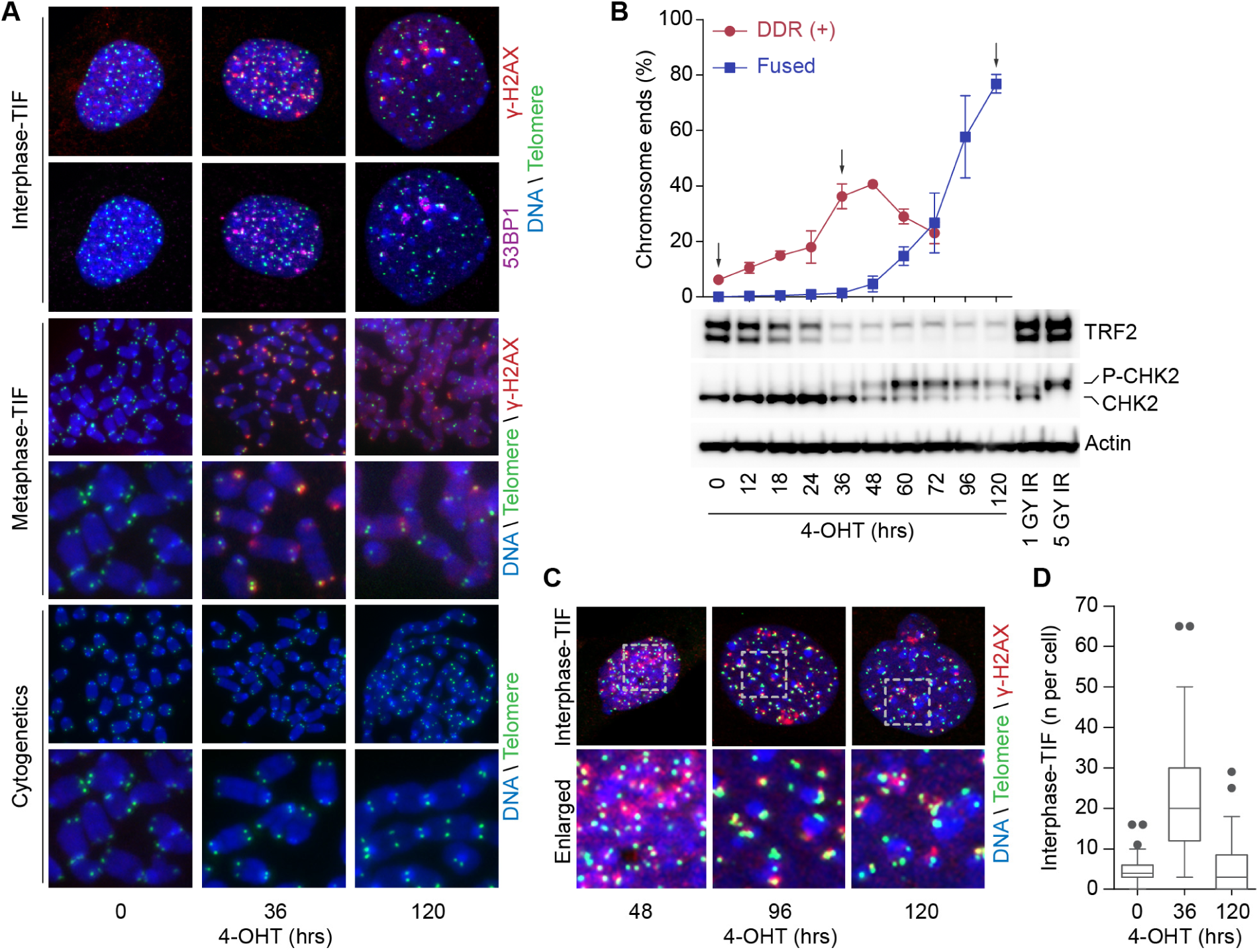
*TRF2* deletion in mouse cells induces a transient DDR-positive/NHEJ-resistant state. A) Examples of interphase-TIF, metaphase-TIF and cytogenetic chromosome assays in *TRF2^F/F^ CreER LgT* MEFs following 4-OHT -induced *TRF2* deletion. B) Quantitation of DDR(+) telomeres by metaphase-TIF assay, and telomere fusions by cytogenetic chromosome spreads, in *TRF2^F/F^ CreER LgT* MEFs following *TRF2* deletion with 4-OHT (mean ± s.e.m., n=3 biological replicates of ≥ 30 metaphase spreads per replicate). Western blots of cell whole extracts from the indicated times are shown. Irradiated samples are positive controls demonstrating phosphorylation of the ATM target CHK2. C) Additional examples of interphase-TIF assays following 4-OHT treatment. D) Quantitation of interphase-TIF in *TRF2F/F CreER LgT* MEFs cells following 4-OHT treatment (three biological replicates scoring ≥ 50 nuclei per replicate are compiled into a Tukey box plot).

A defining feature of intermediate-state telomeres is that the corresponding DDR is distinct from the response induced by genomic double strand breaks. Intermediate-state telomeres in humans activate ATM without downstream CHK2 phosphorylation or G2/M checkpoint activation (Cesare et al., 2013). In agreement with human cells, DDR-positive/fusion resistant telomeres in *TRF2^F/F^ CreER LgT* MEFs following 18-36 hours 4-OHT treatment induced a partial ATM response where CHK2 was not phosphorylated (Figure 1B). CHK2 phosphorylation coincided temporally with the later induction of telomere fusions (Figure 1B). Telomere DDR activation induced by *TRF2* deletion also did not activate the G2/M checkpoint (Figure S2A). Onset of telomere fusions in *TRF2^F/F^ CreER LgT* MEFs corresponded with mitotic arrest and genome duplication events as seen in human cells (Figures S2B-G) (Cesare et al., 2013; Hayashi et al., 2015; Pampalona et al., 2012). However, unlike human cells, the mitotic arrest was shorter in duration and was not lethal (Figures S2E-G). Cumulatively, the data indicate that mouse telomeres exhibited a transient DDR-positive/NHEJ-resistant state following *TRF2* deletion that is phenotypically consistent with intermediate-state telomeres described in human cells.

### Visualizing t-loops in mouse cells

To determine how chromosome end structure relates to the kinetics of telomere deprotection, we used super-resolution microscopy to visualize telomeres from *TRF2^F/F^ CreER LgT* cells at 0, 36 or 120 hours of 4-OHT treatment (Figure S3A) (Doksani et al., 2013). These time points corresponded with protected telomeres (0 h), DDR-positive/NHEJ-resistant telomeres (36 h), and telomere fusions (120 h) (Figure 1B). To maintain t-loops during sample preparation, it is necessary to covalently cross-link duplex DNA within the t-loop junction with trioxsalen and UV light (Doksani et al., 2013; Griffith et al., 1999). After cross-linking, samples were deposited on glass coverslips by cyto-centrifugation in the presence of a mild detergent lysis to decompact the condensed telomeric chromatin, and the telomeres stained by fluorescent *in situ* hybridization (FISH) (Figure S3B) (Doksani et al., 2013).

We successfully visualized murine t-loops with STORM (Figure 2A). However, due to the duration of STORM image capture of up to 30 minutes per field, we developed imaging protocols to visualize t-loops with the more rapid super-resolution imaging modalities of structured illumination (SIM), stimulated emission depletion (STED), and Airyscan microscopy, which typically require 1 to 3 minutes per image. Approximate xy resolution for each of these imaging modalities were: STORM (30 nm), STED (50 nm), SIM (110 nm) and Airyscan (140 nm). Despite reduced resolution, t-loops were effectively imaged using each type of super-resolution microscopy (Figure 2B). Trioxsalen crosslinking efficiency was measured across the experimental time course on bulk genomic DNA, which approximates telomeric DNA crosslinking efficiency (Doksani et al., 2013). For this assay, trioxsalen/UV crosslinked nuclei were micrococcal nuclease digested and the DNA separated on agarose gels. Di-nucleosome bands were isolated, and crosslinking efficiency determined as the percentage of duplex di-nucleosome DNA remaining, relative to input, after denaturation and snap cooling (Figure 2C)(Doksani et al., 2013). No change in cross-linking efficiency was observed across experimental time points, indicating 4-OHT treatment did not impact trioxsalen/UV cross-linking. A previous report demonstrated that telomere DNA crosslinking is not impacted by TRF2 deletion (Doksani et al., 2013).

**Figure 2.**
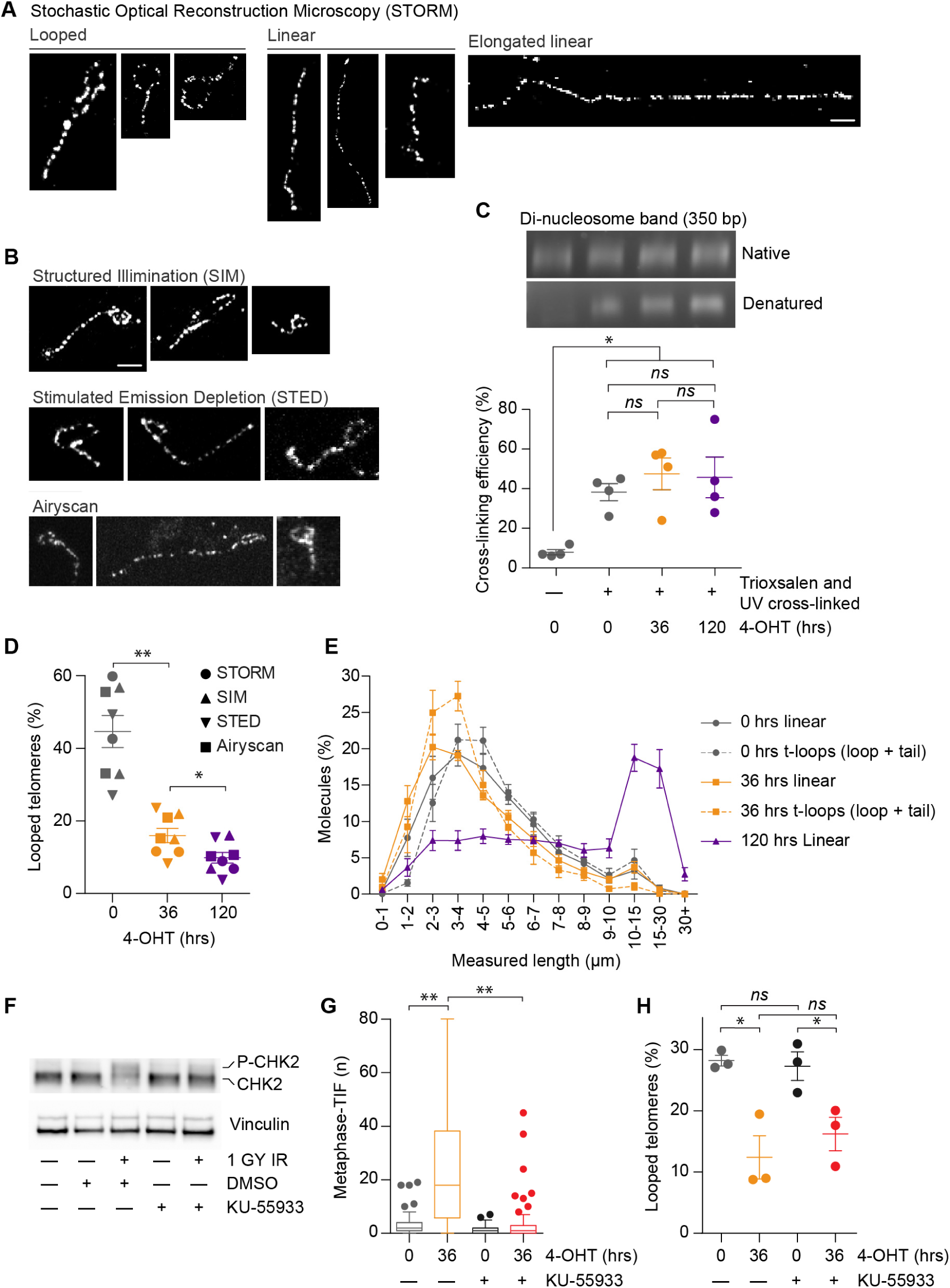
The ATM-dependent telomere DDR occurs with a telomere conformational change from t-loops to linear unfused chromosome ends. A) Examples of telomere structures in *TRF2^F/F^ CreER LgT* MEFs observed with STORM microscopy (bar = 1 µm). B) Examples of t-loops from *TRF2^F/F^ CreER LgT* MEFs observed by SIM, STED and Airyscan microscopy (bar = 1 µm). C) Cross-linking efficiency test for *TRF2F/F CreER LgT* following 4-OHT treatment (mean± s.e.m., n = 4 biological replicates, * *p < 0.05, ns* = not significant, student t-test). D) Percentage of looped telomeres observed by super-resolution microscopy at the indicated times of 4-OHT treatment. Calculations exclude ambiguous molecules from the data set (mean ± s.e.m., n = 8 biological replicates of = 449 molecules per replicate, * *p < 0.05*, ** *p < 0.01*, student t-test). Time points correspond to arrows in (Figure 1B). E) Length distribution of telomere molecules from (D) (mean ± s.e.m., n = 8 biological replicates of = 449 molecules scored per replicate). F) Western blot of whole cell extracts from *TRF2F/F CreER LgT* MEFs following treatment with DMSO or KU-55933 and ionizing radiation. G) Quantitation of DDR(+) telomeres by metaphase-TIF assay in *TRF2F/F CreER LgT* MEFs treated with or without 4-OHT and KU-55933 (n = 3 biological replicates of 30 metaphase spreads per replicate compiled in a Tukey box plot, ** *p < 0.01*, Mann-Whitney test). H) Percentage of looped telomeres observed by Airyscan microscopy at the indicated times of 4-OHT treatment +/-KU-55933. Calculations exclude ambiguous molecules from the data set (mean ± s.e.m., n = 3 biological replicates of = 524 molecules scored per replicate, * *p < 0.05, ns* = not significant, student t-test).

### Telomere DDR activation in MEFs correlates with linearized telomeres

Super-resolution imaging was repeated in eight biologically independent experiments, with each microscopy modality used to capture images for two replicates. Telomeres were visually scored as looped, linear or ambiguous in blinded samples (Figure S3C and Table S1). We observed consistent results between the four imaging modalities, which identified a significant reduction in t-loop frequency corresponding with DDR activation at 36 hours (Figures 2D, S3D, and Table S1). Telomere contour lengths were measured in the super-resolution images. Measurements from the 0 and 36-hour samples revealed similar length distributions of linear and looped (combined measurement of the loop plus the tail) telomeres (Figure 2E). Suggesting that linear structures observed at these time points represent either linear unfused telomeres or t-loops not captured by fixation. In contrast, elongated linear telomere molecules were prevalent at 120 hours, in agreement with cytological visualization of telomere-telomere fusions (Figure 1B, 2A, and 2E).

A conformational change from t-loops to linear telomeres with DDR activation at 36 hours could be the result of: nucleolytic cleavage or degradation of the telomeric DNA and/or 3’ overhang; resolution of t-loops into shortened linear telomeres and telomere circles (t-circles) (Schmutz et al., 2017); or t-loop unfolding resulting in linear telomeres with no change in telomere length or overhang degradation. Separation of telomere restriction fragments on pulsed field gels corresponded with the super-resolution imaging data and verified telomere lengths were unaltered with 36 hours of 4-OHT treatment (Figures 2E, S4A and S4B). Overhang degradation and t-circles were also absent at 36 hours (Figures S4B-D), consistent with t-loop unfolding as the mechanism of telomere linearity occurring with DDR activation.

To determine the relationship between telomere linearization and ATM activation, we repeated t-loop visualization in *TRF2^F/F^ CreER LgT* cells treated with the ATM inhibitor KU-55933 (Hickson et al., 2004). KU-55933 inhibited ATM phosphorylation of its downstream target CHK2 following irradiation in *TRF2^F/F^ CreER LgT* cells (Figure 2F). KU-55933 also suppressed TIF formation at 36 hours of 4-OHT treatment, but did not affect telomere linearization as measured by Airyscan microscopy (Figures 2G, 2H, S4E and Table S1). Telomere linearity at 36 hours, when the DDR is active, thus occurs independently of ATM activation.

### The TRFH domain of TRF2 suppresses ATM and regulates t-loop formation

TRF^cT^ is a separation of function allele containing the TRF2 basic, hinge, and myb domains, and the TRFH domain from TRF1 (Okamoto et al., 2013). The TRFH domain of TRF2 is responsible for homodimerization, ATM inhibition, and alteration of telomere DNA topology (Benarroch-Popivker et al., 2016; Broccoli et al., 1997; Okamoto et al., 2013). Deleting *TRF2* in TRFcT expressing cells results in DDR-positive but NHEJ-resistant telomeres due to retention of the iDDR motif in the TRF2 hinge domain (Okamoto et al., 2013). We transduced Myc-TRF2 or Myc-TRFcT in *TRF2^F/F^ CreER LgT* cells and selected for transgene expression. Endogenous *TRF2* was deleted with 4-OHT for 120 hours and the corresponding telomere phenotypes examined (Figures 3A-C). Expression of Myc-TRF2 rescued both metaphase-TIF and telomere fusions in *TRF2F/F CreER LgT* cells after 120 hours 4-OHT treatment, whereas TRF^cT^ expression led to abundant metaphase-TIF but suppressed telomere fusions (Figures 3D-F). Airyscan super-resolution microscopy revealed no reduction in t-loop abundance in Myc-TRF2 over-expressing cells at 120 hours 4-OHT when the DDR remained suppressed. However, t-loop abundance was significantly reduced in Myc-TRFcT cells at 120 hours of 4-OHT, when the telomere DDR was active (Figures 3G-I and Table S1). These observations indicate that the TRFH domain of TRF2 regulates both ATM inhibition and t-loop formation. Additionally, these data identify that linear DDR-positive telomeres remain fusion resistant in the presence of a functional TRF2 iDDR motif.

**Figure 3.**
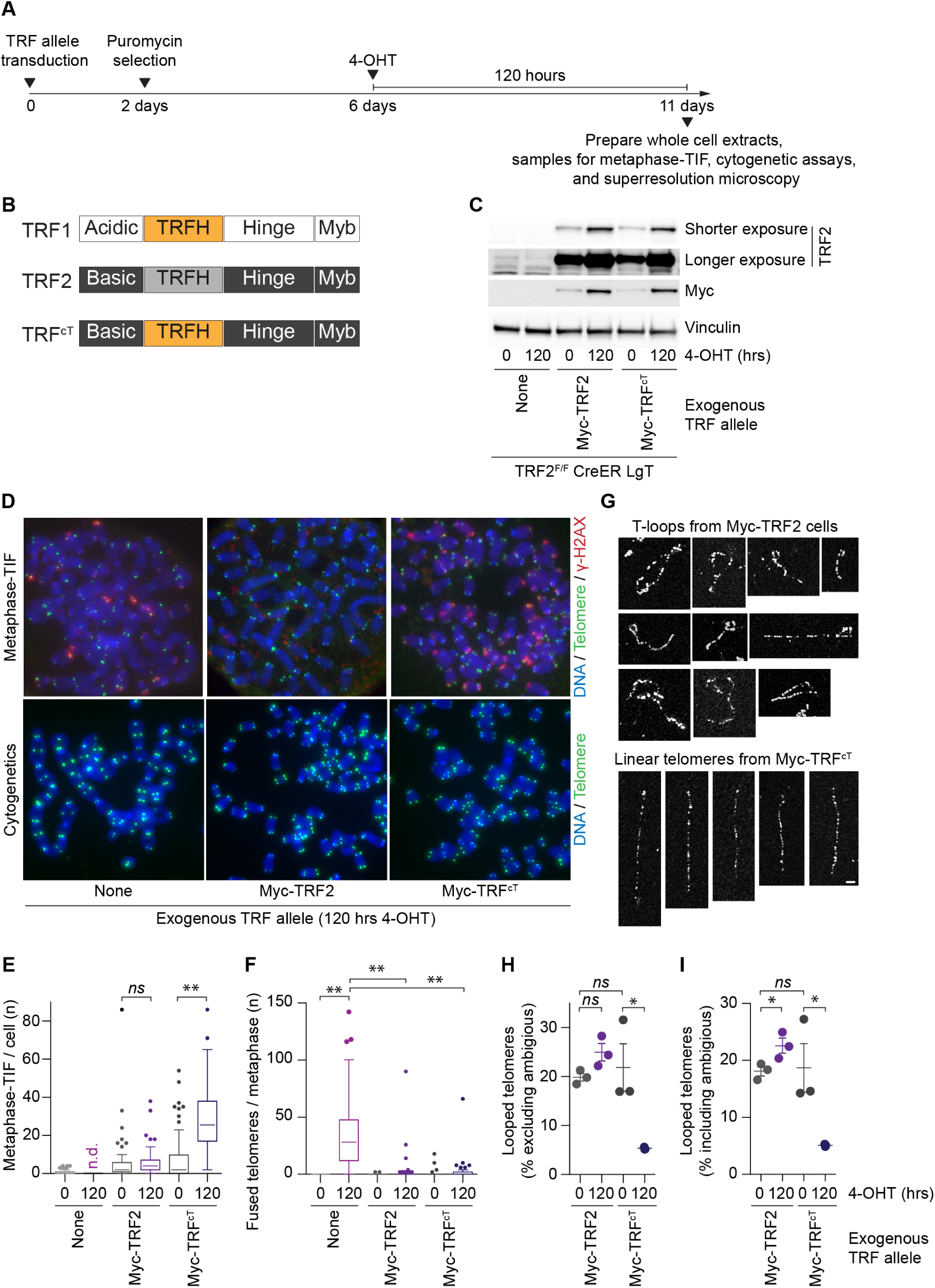
T-loop formation is mediated by the TRFH domain of TRF2. A) Timeline of the experimentation in (C-I) below. B) Graphical depiction of TRF1, TRF2 and TRF^cT^ domains. C) Western blots of whole cell extracts from *TRF2^F/F^ CreER LgT* cells with or without TRF allele over-expression at 0 or 120 hours after 4-OHT treatment. D) Representative metaphase-TIF and cytogenetic chromosome assays from *TRF2^F/F^ CreER LgT* cells with or without Myc-TRF2 or Myc-TRF^cT^ expression, with 120 hours of 4-OHT treatment. E, F) Quantitation of metaphase-TIF (E) and telomere fusions in cytogenetic assays (F) from the experiments represented in (D) (n = 3 biological replicates of 30 metaphase spreads per replicate, compiled in a Tukey box plot, ** *p < 0.01*, Mann-Whitney test, *ns* = not significant, n.d. = not determined). G) Representative t-loops from *TRF2^F/F^ CreER LgT*, and linear telomeres from *TRF2^F/F^ CreER LgT* Myc-TRFcT, after 120 hours 4-OHT (bar = 1 µm). H, I), Percentage of looped telomeres observed by Airyscan microscopy in *TRF2^F/F^ CreER LgT* Myc-TRF2 and *TRF2^F/F^ CreER LgT* Myc-TRFcT, at the indicated times of 4-OHT treatment. Calculations exclusive (H)(mean ± s.e.m., n = 3 biological replicates of ≥ 713 molecules scored per replicate, * *p < 0.05, ns* = not significant, student t-test), and inclusive (I)(mean ± s.e.m., n = 3 biological replicates of ≥ 827 molecules scored per replicate) of ambiguous molecules from the data set are shown. * *p < 0.05, ns* = not significant, student t-test.

### Visualization of t-loops from human cells

Human telomeres are shorter than laboratory mouse telomeres, and chromosome end structures in the eroded telomeres of aged primary human cells cannot be resolved by super-resolution microscopy. Additionally, shorter human telomeres photo-bleach rapidly with STORM and STED imaging. We overcame these biological and technical limitations, and successfully imaged t-loops using Airyscan microcopy in the human HeLa Long Telomere (LT) subclone (Figure 4A). HeLa LT harbors average telomere lengths of approximately 20 kb (O’Sullivan et al., 2014) and while Airyscan has a reduced resolution, it captures images rapidly to minimize photo-bleaching.

**Figure 4.**
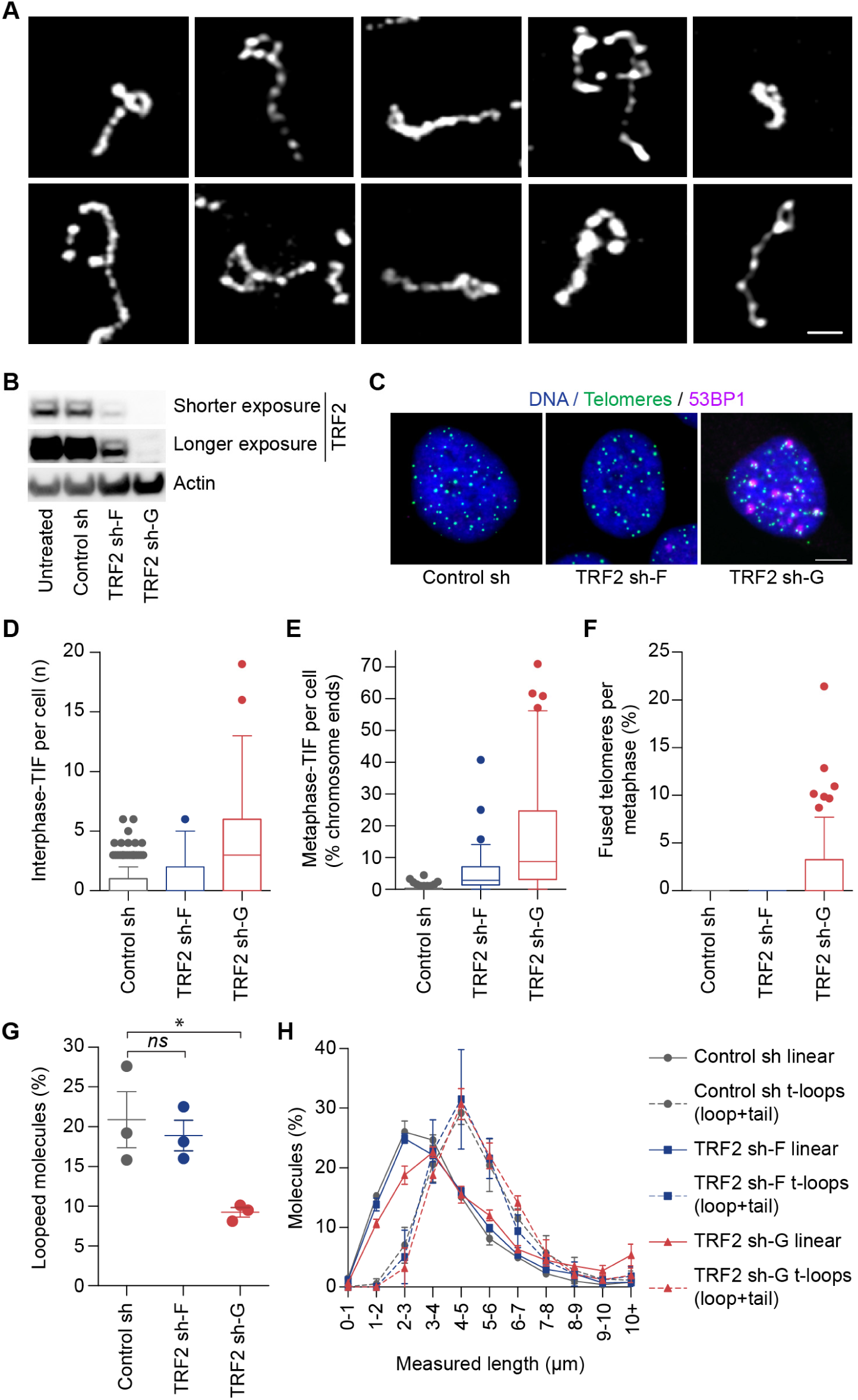
Linear telomeres in human cells occur with induction of the ATM-dependent telomere DDR not simply reduced TRF2. A) T-loops from HeLa LT cells visualized by Airyscan microscopy (bar = 1 µm). B) Western blots of whole cell extracts from HeLa LT cells transduced with TRF2 shRNAs. C) Examples of interphase-TIF assays in shRNA transduced HeLa LT cells. D-F) Quantitation of interphase-TIF (D), metaphase-TIF (E) and telomere fusions (F) in TRF2 shRNA transduced HeLa LT cells [three biological replicates scoring ≥ 80 nuclei (D), or ≥ 30 metaphase spreads (E, F), per replicate are compiled into a Tukey box plot]. G) Quantitation of looped telomeres from HeLa LT observed by Airyscan microscopy. Calculations exclude ambiguous molecules from the data set (mean ± s.e.m., n = 3 biological replicates scoring ≥ 606 molecules per replicate, * *p < 0.05, ns* = not significant, student t-test). h) Length distribution of the telomere molecules scored in (G) (mean ± s.e.m., n = 3 biological replicates scoring ≥ 606 molecules per replicate).

### The telomere DDR in human cells occurs with reduced TRF2 and linear telomeres

Unlike *TRF2* deletion in MEFs, where telomeres inexorably progress towards fusion due to progressive TRF2 turnover, stable TRF2 depletion in human cells induces steady-state levels of ATM-dependent intermediate-state telomeres independent of telomere length (Cesare et al., 2013). The resulting telomere deprotection response is phenotypically consistent with the canonical telomere DDR activated during cellular aging (Cesare et al., 2013). Depending on shRNA effectiveness and cell line, TRF2 depletion may induce no phenotype, a small number of intermediate-state telomeres, or an elevated number of intermediate-state telomeres and some telomere fusions (Cesare et al., 2013). Using lentivector TRF2 shRNAs of different efficiencies (Cesare et al., 2013), we depleted TRF2 in HeLa LT cells (Figures 4B and S5A-D). The resulting phenotypes in HeLa LT were milder than previously observed in HT1080 6TG cells, potentially as a result of different levels of endogenous TRF2 (Figures S5B-D). In HeLa LT, TRF2 sh-F induced a very modest TIF phenotype with no telomere fusions, while TRF2 sh-G induced a more robust TIF phenotype and a low number of telomere fusions (Figures 4B-F, and S5B-D).

Macromolecular telomere structure was visualized with Airyscan microscopy in three independent biological replicates of HeLa LT control shRNA and TRF2 depleted cells (Figure S5E). Measurement of trioxsalen cross-linking efficiency indicated no change between control shRNA and TRF2 shRNA cells (Figure S5F). Interestingly, we observed no reduction in t-loops from HeLa LT TRF2 sh-F cells where the telomere DDR was largely absent, while t-loop prevalence was reduced in HeLa LT TRF2 sh-G cells with an elevated number of intermediate-state telomeres (Figures 4G, S5G, and Table S2). Together, the data indicate that linear telomeres in human cells correlate specifically with activation of the ATM-dependent telomere DDR (TRF2 sh-G), and not simply from reduced TRF2 protein levels (TRF2 sh-F).

Distribution of telomere contour measurements revealed a striking overlap in the length of looped telomeres (combined measurement of the loop plus the tail) from all conditions (Figure 4H). TRF2 sh-G cells exhibited a slight increase in average linear telomere length and more linear molecules of ≥ 10 µm, consistent with the small number of telomere fusions observed in these cells (Figure 4H). In all conditions, the length distributions of looped and linear telomeres overlapped, with the distribution of looped telomeres shifted towards a longer average telomere length (Figure 4H). We expect this discrepancy reflects the experimental limits of visualizing shorter human telomeres. Classifying a molecule as a t-loop requires resolution of an open loop structure. Shorter molecules with small loops are more likely to be misclassified. Additionally, if t-loops do not decondense sufficiently during sample preparation, or if t-loops deposit on slides in a way that masks the loop portion of the molecule, they will be misclassified as linear or ambiguous. These factors are less likely to impact linear molecules, which are more effectively scored regardless of length. Molecules scored as t-loops in human cells likely disproportionally represent longer telomeres in the population that are more efficiently presented in micrographs. For these reasons, we also anticipate the reported percentage of observed t-loops in our experiments underestimates the number of t-loops in the sample population.

### Aurora B regulates telomere linearization corresponding with the mitotic telomere DDR

The non-canonical telomere DDR induced during mitotic arrest is ATM-dependent, and is regulated through an Aurora B kinase-dependent mechanism where TRF2 is partially removed from chromosome ends (Hayashi et al., 2012). We therefore set out to determine the relationship of macromolecular telomere structure to mitotic telomere deprotection.

To perform these experiments, we required HeLa LT cells with a sufficient dynamic range of induced telomere deprotection during mitotic arrest, to observe potential corresponding changes in telomere structure with super-resolution microscopy. HeLa LT cultures enter mitosis 8-10 hours following synchronization and release from a double thymidine block (Figures S6A and 6B).

Synchronizing HeLa LT with a double thymidine block, releasing, and adding colcemid six hours later, induced subsequent mitotic arrest and accumulation of mitotic cells (Figures S6-C). To increase the dynamic range of DDR-positive chromosome ends induced during mitotic arrest, we tested if partial TRF2 depletion sensitized HeLa LT telomeres to mitotic telomere deprotection as shown previously in other cell types (Cesare et al., 2013). We quantified metaphase-TIF in HeLa LT control and TRF2 shRNA cells, following a double thymidine block, release, and colcemid treatment. The conditions that provided the greatest dynamic range of mitotic arrest-induced telomere deprotection, relative to cells without mitotic arrest (8 hours after release), were TRF2 sh-G cultures 18 hours post release (Figure S6D).

To visualize how mitotic telomere deprotection influences telomere macromolecular structure, HeLa LT TRF2 sh-G cells were synchronized with a double thymidine block and released. At the time of release, the cell culture had been selected after shRNA transduction for 5 days. Colcemid was added to the cultures six hours later to induce a subsequent mitotic arrest (Figure 5A). Six hours after release, the cultures were also treated with vehicle (DMSO) or Hesperadin. Hesperadin is an Aurora B inhibitor previously shown to suppress mitotic telomere deprotection (Hauf et al., 2003; Hayashi et al., 2012). Importantly, Hesperadin treatment does not inhibit mitotic ATM activity in response to genomic double strand breaks, signifying that Aurora B inhibition prevents mitotic telomere deprotection by modulating telomere function (Hayashi et al., 2012). Mitotic cells were collected by shake-off 18 hours after release from the double thymidine block, and samples were prepared for metaphase-TIF assays, super-resolution microscopy, and to determine trioxsalen cross-linking efficiency (Figure 5A).

**Figure 5.**
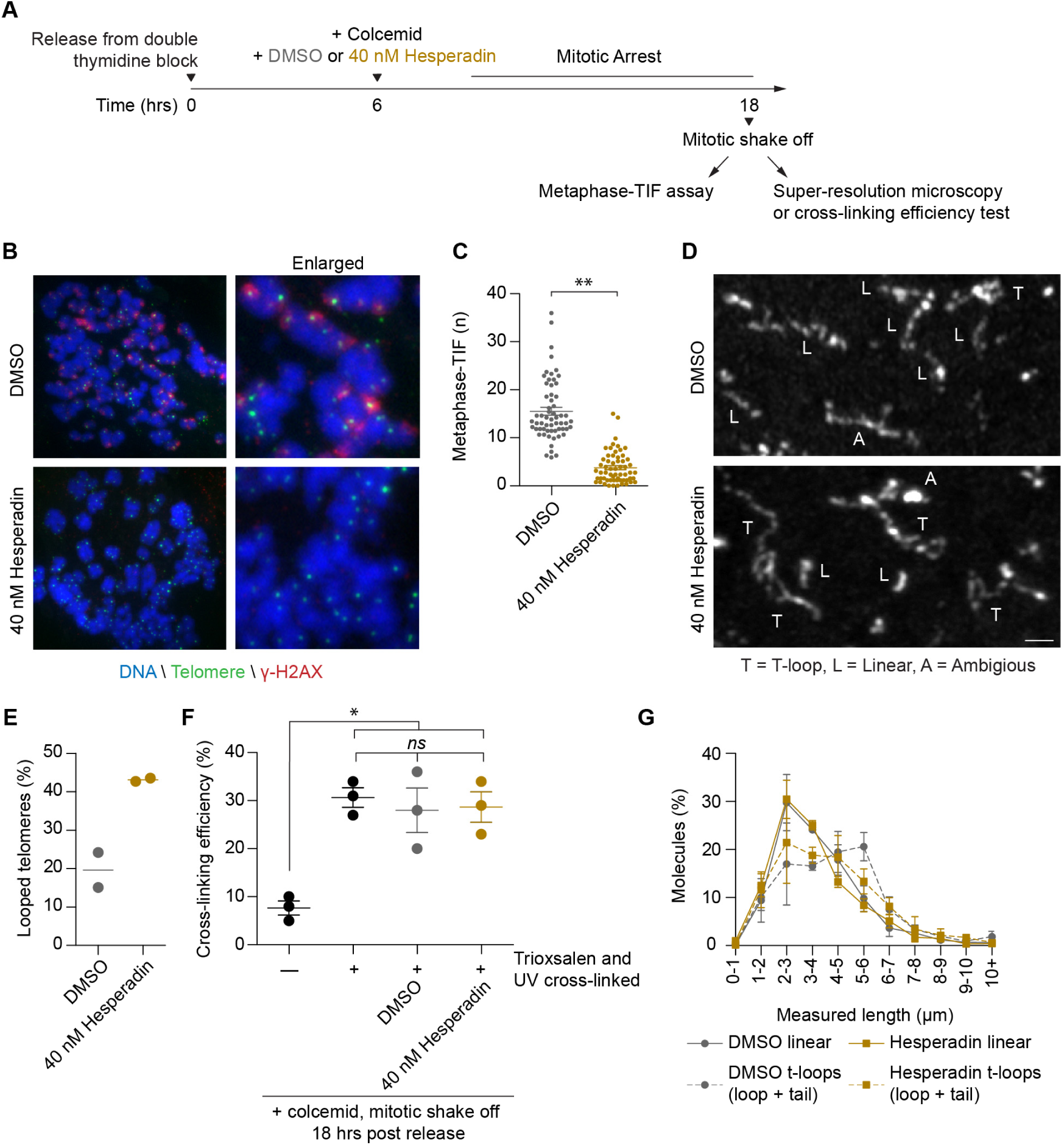
Mitotic arrest induces an Aurora B-dependent telomere conformational change from looped to linear coinciding with telomere DDR activation. A) Timeline of the experimentation in (B-G) below. B) Examples of metaphase-TIF assays in HeLa LT TRF2 sh-G cells following mitotic arrest in the presence of Hesperadin or vehicle. C) Quantitation of the metaphase-TIF assays shown in (B) (mean ± s.e.m., all data points from two biological replicates of at least 30 metaphases). D) Examples of Airyscan microscopy visualization of telomere structure in HeLa LT TRF2 sh-G cells following mitotic arrest in the presence of Hesperadin or vehicle. E) Quantitation of looped telomeres in mitotically arrested HeLa LT cells. Calculations exclude ambiguous molecules from the data set (n = 2 biological replicates of = 794 molecules per replicate, line depicts the mean). F) Determination of cross-linking efficiency in mitotically arrested HeLa LT cells treated with DMSO or Hesperadin (mean ± s.e.m., n = 3 biological replicates, * *p < 0.05, ns* = not significant, student t-test). G) Length distribution of the telomere molecules scored in (E) (mean ± range, n = two biological replicates scoring = 794 molecules per replicate).

Metaphase-TIF assays confirmed that mitotic-arrest induced telomere deprotection was suppressed with Hesperadin treatment, but was abundant in the vehicle treated cells (Figures 5B and 5C). In agreement, t-loops were prevalent in Hesperadin treated cultures with a greater percentage of protected telomeres, and reduced in the vehicle treated cells with abundant mitotic telomere deprotection (Figures 5D, 5E, S6E and Table S2). Mitotic arrest, Hesperadin or DMSO did not impact trioxsalen cross-linking efficiency (Figure 5F). Telomere length contour measurements revealed an overlap in the lengths of linear and looped telomeres (combined measurement of the loop plus the tail) isolated from all conditions, with a skew towards a longer average length of looped telomeres as described in the above HeLa LT experiments (Figure 5G). Together, the data are consistent with a mechanism where Aurora B regulates an active telomere conformational change from looped to linear during mitotic arrest coincident with ATM-dependent telomere DDR activation.

### Intermediate-state telomeres are resistant to NHEJ

Data presented above show increased linear telomeres corresponding with telomere DDR activation, but not telomere fusions, in *TRF2^F/F^ CreER LgT* MEFs at 36 hours 4-OHT (Figure 2), TRF^cT^ expressing *TRF2* null MEFs (Figure 3), and HeLa LT TRF2 sh-G cells (Figure 4): implying separate regulation of ATM and NHEJ at chromosome ends. It is possible, however, that cell cycle regulation of DNA repair leads to the misconception that ATM and NHEJ are independently controlled. Telomere fusion requires migration and co-localization of DDR-positive telomeres in G1-phase when NHEJ is active (Dimitrova et al., 2008; Lottersberger et al., 2015). This is anticipated to take longer than synapsis of DNA ends at a double strand break, which remain in immediate proximity after damage induction. There may be insufficient time in cycling cells for DDR-positive telomeres to migrate, synapse and fuse during G1-phase.

To test this directly, we depleted TRF2 in IMR90 E6E7 fibroblasts and arrested cells in G1-phase (Figure 6A and 6B). In IMR90 E6E7, TRF2 sh-F induces intermediate-state telomeres, while TRF2 sh-G induces both intermediate-state telomeres and telomere fusions (Cesare et al., 2013). IMR90 E6E7 control shRNA and TRF2 shRNA cells were arrested in G1-phase with Lovastatin treatment for 24 hours before release with mevalonic acid (Figure 6A and 6C) (Javanmoghadam-Kamrani and Keyomarsi, 2008). A population of cells entered S-phase 16 hours after release and progressed through mitosis around 24 hours (Figure 6C). In IMR90 E6E7 TRF2 shRNA cells, interphase-TIF were present in the synchronized G1 population, and at all time points following release (Figures 6D and 6E). The number of interphase TIF remained at steady state levels throughout the experiment. Because the telomere DDR is resolved following telomere fusion, persistent interphase-TIF are consistent with intermediate-state telomeres. Notably, irradiation of Lovastatin treated IMR90 E6E7 cells induced an increase and subsequent reduction in γ-H2AX signal intensity, consistent with induction and repair of double-strand breaks (Figure S7). Lovastatin induced G1-arrest therefore did not impact double-strand break repair.

**Figure 6.**
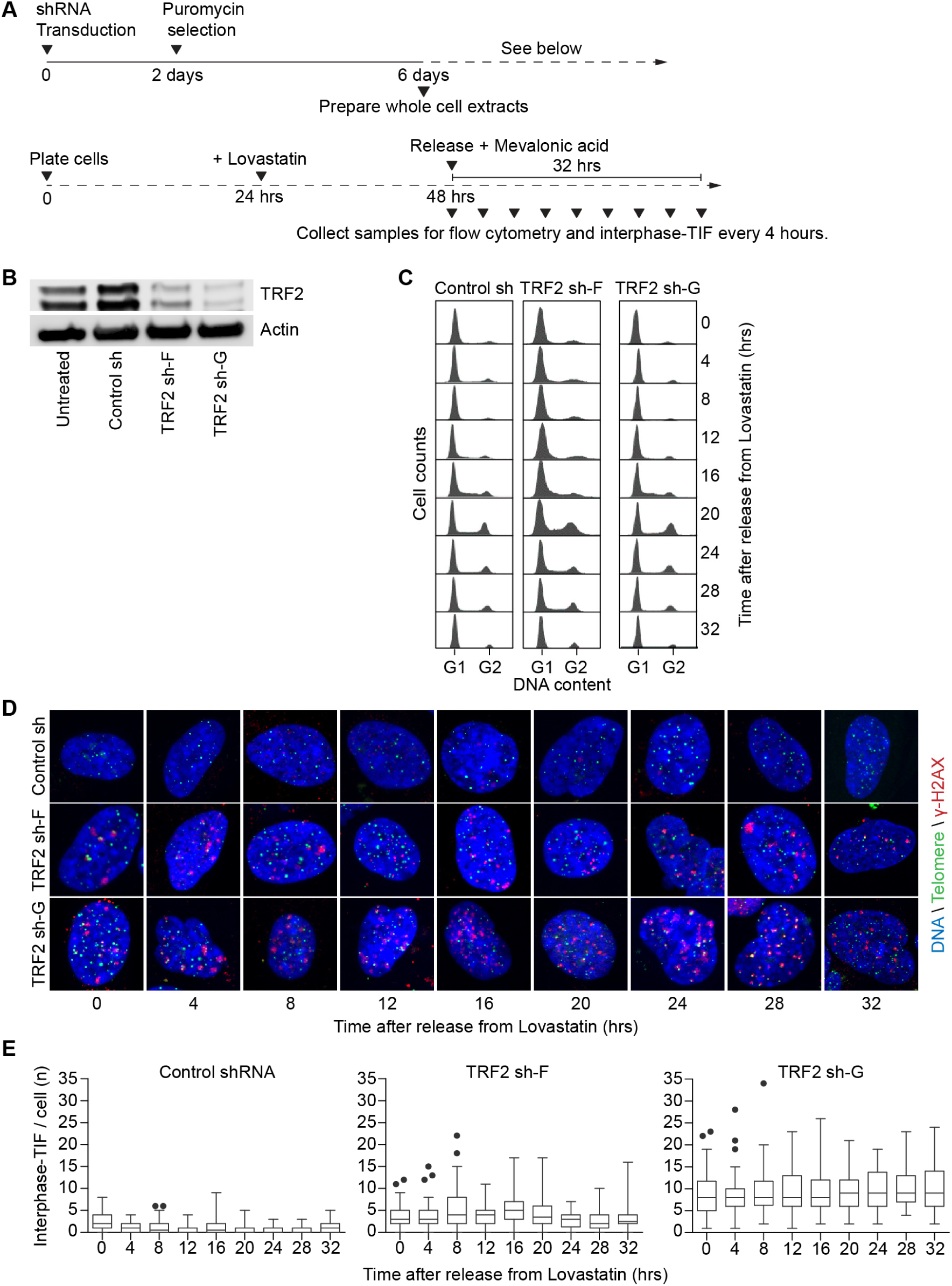
Interphase-TIF persist during an extended G1-phase arrest and the following cell cycle. A) Timeline of the experimentation in (B-E) below. B) Western blots on whole cell extracts from IMR90 E6E7 cells transduced with the indicated shRNAs. C) Cell cycle distribution of IMR90 E6E7 shRNA transduced cultures after 24 hours of Lovastatin treatment and release. D) Examples of interphase-TIF assays in shRNA transduced IMR90 E6E7 cultures following 24 hours of Lovastatin treatment and release. E) Quantitation of interphase-TIF from the experiment in (D) (n = 3 biological replicates scoring ≥ 30 nuclei per replicate are compiled into a Tukey box plot).

Quantitative metaphase-TIF analysis performed 24 hours after release from Lovastatin, revealed the extended G1-arrest did not impact progression of DDR-positive telomeres through the following cell division (Figures 7A-C). Strikingly, prolonged G1-arrest also did not induce telomere fusions in IMR90 E6E7 TRF2 sh-F cells, nor did it increase the number of telomere fusions in IMR90 E6E7 TRF sh-G cells (Figure 7D, and 7E). Cumulatively, these data confirm intermediate-state telomeres are fusion resistant, even during an extended G1-phase when telomeres are provided ample opportunity to synapse and fuse.

**Figure 7.**
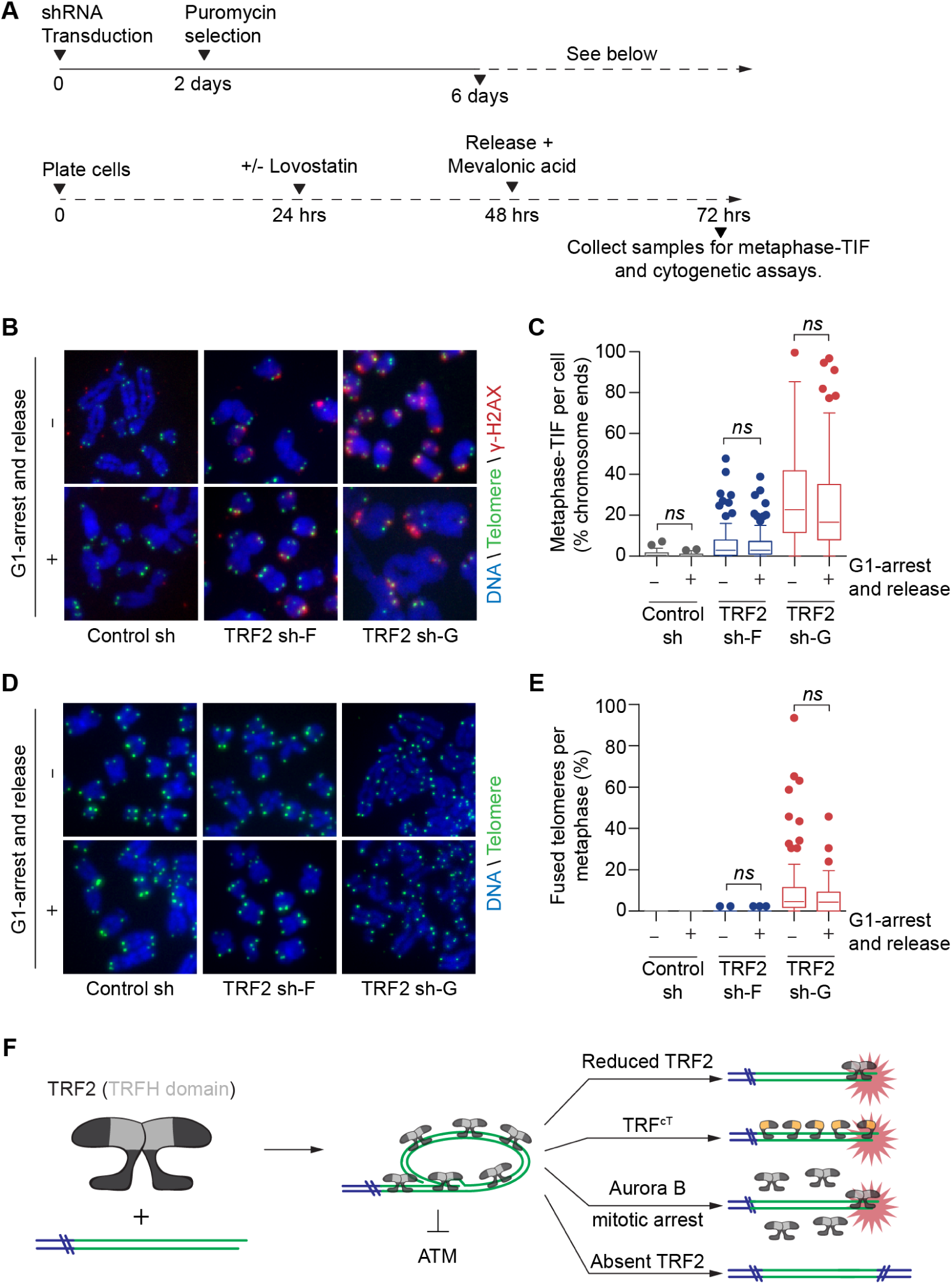
Intermediate-state telomeres do not fuse during an extended G1-arrest. A) Timeline of the experiments in (B-E) below. B) Examples of Metaphase-TIF assays from shRNA transduced IMR90 E6E7 cells following G1-arrest and release. C) Quantitation of metaphase-TIF from (B) (n = 3 biological replicates scoring ≥ 30 mitotic cells per replicate are compiled into a Tukey box plot, *ns =* not significant, Mann-Whitney test). D) Examples of cytogenetic chromosome spreads from shRNA transduced IMR90 E6E7s following G1-arrest and release. E) Quantitation of telomere fusions from (D) (n = 3 biological replicates scoring ≥ 30 mitotic spreads per replicate are compiled into a Tukey box plot, *ns =* not significant, Mann-Whitney test). G) Graphical model depicting t-loops regulating ATM activity. The telomere DDR is represented by a starburst.

## DISCUSSION

Here we present the first comprehensive study examining the role of telomere structure through the spectrum of telomere deprotection. To achieve this, we developed rapid methods to visualize murine telomere structure using STED, SIM and Airyscan microscopy, which also enabled effective visualization of human t-loops with Airyscan. Our results are consistent with independent telomere regulation of ATM and NHEJ activity, and suggest t-loops function specifically to regulate DDR activation at chromosome ends. These data define the molecular identity of intermediate-state telomeres and provide insight into mechanisms of telomere-dependent tumor suppression (Figure 7F).

### Telomere macromolecule structure relative to telomere protection

We visualized telomere structure with four different types of super-resolution microscopy. Despite their lower resolution, SIM, STED and Airyscan resolved t-loops effectively, and provided increased experimental output through faster data capture. Reduced photobleaching with Airyscan also allowed us to image the elongated telomeres in HeLa LT cells. Collectively, we observed experimental means of 19.6 to 44.6% looped telomeres in cells with wild type shelterin, consistent with previous studies (Doksani et al., 2013; Griffith et al., 1999).

Experimental and imaging limitations contributing to underrepresentation of t-loops in electron or super-resolution micrographs are described above, and in detail elsewhere (Doksani et al., 2013; Griffith et al., 1999). Cross-linking efficiency also impacts observed t-loop frequency. We found cross-linking efficiencies in ∼350 bp dinucleosome DNA ranged from 25 to 75% in MEFs (mean ± s.d., 43.8 ± 14.9) and 18 to 48% in HeLa LT (29.3 ± 7.8). The structure of t-loop junctions and extent of duplex DNA created by overhang strand invasion remains unclear; however, extrapolation of our cross-linking data suggest a hypothetical t-loop junction consisting of 350 bp duplex DNA would be cross-linked in approximately 30 to 45% of occurrences. This roughly approximates the t-loop percentages we observed in cells with protected telomeres. In all experiments we observed a concomitant reduction in t-loops coinciding with ATM activation at chromosome ends, despite no change in cross-linking efficiency. Biochemical data indicated telomere linearity with DDR activation proceeds by t-loop “unfolding", as we observed no evidence of duplex telomeric DNA and/or 3’-overhang degradation, nor t-circle formation.

Linear telomeric DNA observed at 120 hrs of 4-OHT treatment in *TRF2^F/F^ CreER LgT* MEFs was elongated relative to 0 hrs, consistent with the preponderance of telomere-telomere fusions at this time point. In contrast, linear telomeres coinciding with DDR activation in human and mouse cells, all overlapped in length distribution with looped telomeres from the respective negative control with protected chromosome ends, indicating the linear DDR-positive telomeres had not fused. Cumulatively, our observations are consistent with ATM activation at telomeres coinciding specifically with a structural change from t-loops to unfused linear chromosome ends. Telomere fusion in *TRF2^F/F^ CreER LgT* MEFs occurred thereafter in a temporally separate event.

### A putative role for t-loops in chromosome end protection

Through expression of TRF^cT^ in a TRF2 null background, we identified that the TRFH domain of TRF2 functions to both suppress ATM activity and modulate t-loop formation: indicating that rescue of DDR activity coincided specifically with the ability of TRF2 to form t-loops. Additionally, we found ATM-dependent telomere DDR activation in HeLa LT cells occurred not as a consequence of reduced TRF2 per se, but specifically with t-loop unfolding resulting in telomere linearity. We also find that t-loop unfolding occurs independent of ATM activity, indicating telomere linearity was not the outcome of DDR activation. This suggests instead that telomere linearity precedes ATM activation at chromosome ends. In data not shown, we tried exhaustively to visualize telomere structure in the context of DDR activation using super-resolution microcopy, but were unable to identify fixation conditions enabling us to resolve telomere structure while retaining protein targets of ATM activation. However, it is well established that DNA ends, single-strand/double strand DNA junctions in particular, are potent activators of the ATM-dependent DDR (Lee and Paull, 2005; Shiotani and Zou, 2009). Presentation of the naturally occurring DNA end, specifically the single-strand/double-strand junction provided by the 3’-telomere overhang, is likely an ideal ATM activating substrate. We propose the most logical interpretation of our data, is that t-loops function specifically to suppress ATM activation through sequestration of chromosome ends, independent of telomere protection against NHEJ.

Our observations of the telomere DDR in 4-OHT treated *TRF2^F/F^ CreER LgT* MEFs, with or without TRF^cT^ expression, are consistent with intermediate-state telomeres in human cells. Specifically, DDR-positive/fusion-resistant telomeres were observed in all cell cycle phases, arising from both leading and lagging-strand replicated telomeres, and in some circumstances occurring on a preponderance of chromosome ends (Cesare et al., 2013; Cesare et al., 2009; Hayashi et al., 2012; Kaul et al., 2012). The simplest interpretation is that all mammalian telomeres suppress ATM-activation through a common mechanism. We are unable to determine the precise percentage of chromosome ends that are arranged into t-loops, but propose that conceptualizing telomere protection from a t-loop perspective synthesizes data generated from multiple laboratories over time.

### T-loops as conformational switches that regulate ATM activity at chromosome ends

We suggest telomeres function like molecular switches that can be reversibly shifted between stable states. In this model, the TRFH domain of TRF2 regulates ATM through a “conformational switch”. When telomeres are in a t-loop, the chromosome end is hidden, preventing ATM activation. In this conformation the switch is “off” and the telomere DDR is suppressed. Telomere linearization exposes the chromosome end as an ATM substrate, turning the switch “on” and activating the DDR. NHEJ remains suppressed at these telomeres, but is dependent on sufficient TRF2 containing a functional iDDR motif being bound to the DDR-positive chromosome end. Consistent with a reversible switch, the ATM-dependent telomere DDR can be turned on and off without accompanying telomere fusion, suggesting linear unfused telomeres can reform t-loops (Cesare et al., 2013; Kaul et al., 2012; Konishi and de Lange, 2008). A t-loop model of ATM regulation is also consistent with TRF2 controlling ATM by a telomere structure dependent upon telomere length (Cesare et al., 2009; Karlseder et al., 2002; Kaul et al., 2012), alteration of telomere DNA topology (Benarroch-Popivker et al., 2016), and sufficient TRF2 binding (Cesare et al., 2013). Shortfalls in any of these categories would be expected to correspond with impaired t-loop formation or maintenance, thereby changing the telomere conformation to linear, and switching ATM activation on.

### Potential role for t-loops in regulating cell outcomes

Technical limitations prevented us from visualizing telomere structure in aged primary cells with shortened telomeres. However, the phenotypic similarity of induced intermediate-state telomeres, and those that arise during cellular aging, suggest the telomere DDR may be initiated from the same underlying mechanism. In the future, it will be interesting to determine if telomere erosion impairs t-loop formation leading to telomere linearity and DDR activation as a means of regulating cellular aging and telomere-dependent tumor suppression. We were, however, able to directly demonstrate that Aurora B kinase regulates telomere linearization during mitotic arrest. The mitotic arrest-dependent telomere DDR at cell crisis, or in cells treated with chemotherapeutic mitotic poisons, contributes to mitotic cell death (Hayashi et al., 2015). Aurora B kinase thus functions to actively alter telomere structure during mitotic arrest to impact cell outcomes, suggesting t-loops function as a conduit to signal cell death in response to mitotic abnormality. Cumulatively, the results of this study demonstrate that macromolecular structure adds a degree of regulation to the protein and DNA composition of telomeres, thus enabling telomeres to regulate different outcomes in the DDR and DNA repair pathways and facilitate growth arrest without threatening genome stability.

## ACKNOWLEDGEMENTS

All data are archived at the Children’s Medical Research Institute (CMRI). We thank Scott Page and the CMRI ACRF Telomere Analysis Centre supported by the Australian Cancer Research Foundation, the Australian Centre for Microscopy & Microanalysis at the University of Sydney, the Microbial Imaging Facility at the University of Technology Sydney, and the University of New South Wales Biomedical Imaging Facility, for use of microscopy infrastructure. Leszek Lisowski is thanked for producing lentiviral vectors and Eros Lazzerini Denchi for providing critical reagents. Tracy Bryan, Roger Reddel, and members of Bryan, Reddel, Pickett and Cesare laboratories are thanked for their comments. H.A.P. is supported by the Cancer Council NSW (RG 16-09). K.G. is supported by the Australian Research Council (CE140100011, LP140100967, LE150100163) and the National Health and Medical Research Council of Australia (1059278, and 1037320). A.J.C. is supported by grants from the National Health and Medical Research Council of Australia (1053195, 1106241, 1104461), the Cancer Council NSW (RG 15-12), and the Cancer Institute NSW (11/FRL/5-02).

## AUTHOR CONTRIBUTIONS

A.J.C., D.V.L., and K.G. conceived the study; D.V.L., R.R.J.L., S.F, K.G. and A.J.C. designed experiments; D.V.L., R.R.J.L., S.F, T.A.B., G.R.F., H.A.P., and A.J.C. conducted experimentation and analyzed data; A.J.C. and D.V.L. wrote the manuscript with editorial assistance from R.R.J.L., S.F, G.R.K., H.A.P., and K.G.

## DECLARATION OF INTERESTS

The authors declare no competing interests.

## METHODS

### Cell culture

*TRF2^Floxed/Floxed^ Rosa26-CreERT2 pBabeSV40LT* MEFs were a gift from Eros Lazzerini-Denchi (The Scripps Research Institute, La Jolla, CA, USA)(Okamoto et al., 2013), HT1080 6TG cells were a gift from Eric Stanbridge (University of California at Irvine, USA) and IMR90 E6E7, HeLa LT and U-2OS cells were a gift from Jan Karlseder (The Salk Institute, La Jolla, CA, USA). The identity of all cell lines was verified by Cell Bank Australia using short tandem repeat profiling. MEFs, HT1080 6TG, HeLa LT and U-2OS cells were grown at 37°C, 10% CO2 and atmospheric oxygen in DMEM (Life Technologies) supplemented with 1% non-essential amino acids (Life Technologies), 1% Glutamax (Life Technologies), and 10% bovine growth serum (Hyclone). IMR90 E6E7 cells were cultured at 37°C, 10% CO2 and 3% O2 in DMEM supplemented with 1% non-essential amino acids, 1% Glutamax, and 10% fetal bovine serum (Life Technologies). Cells were tested for mycoplasma contamination (MycoAlert, LT07-118, Lonza) and were all found to be negative. *TRF2* was deleted in *TRF2F/F CreER LgT* cells by adding 1 µM 4-hydroxytamoxifen (Sigma-Aldrich) to the culture media. Aurora B was inhibited with Hesperadin (Selleck chemicals) and ATM with KU-55933 (Calbiochem).

### Myc-TRF2 and Myc-TRF^cT^ transduction

Retroviral vectors were created by transfecting 293T cells with pVPack-GP and pVPack-Ampho (Ailent Technologies), along with pLPC-Myc-TRF2 (a gift from Titia de Lange, Addgene plasmid #16066) or pLPC-Myc-TRFcT (a gift from Eros Lazzerini Denchi, Addgene plasmid #44573). Viral containing supernatants were used to infect *TRF2Floxed/Floxed Rosa26-CreERT2 pBabeSV40LT* MEFs and stably transfected cells selected with 1 µg/ml puromycin.

### TRF2 shRNA transduction

High titre, purified pLKO.1 derived lentiviral vectors, harboring a non-targeting control shRNA (a gift from David Sabatini, Addgene plasmid #1864), TRF2 sh-F (Open Biosystems, TRCN0000004811), or TRF2 sh-G (Open Biosystems, TRCN0000018358), were created in the Salk Institute Gene Transfer, Targeting and Therapeutics core (La Jolla, CA, USA) or the CMRI Vector and Genome Engineering Facility as described elsewhere (Crabbe et al., 2012; Hayashi et al., 2012). Cell cultures were transduced with an MOI of 10 for 48 hours, then selected in normal growth media supplemented with 1 µg/ml Puromycin.

### Cell synchronization and cell cycle arrest

HeLa LT cells were synchronized in G1/S using a double thymidine block, and colcemid was added to 100 ng/ml to induce mitotic arrest, as described elsewhere (Hayashi et al., 2012). IMR90 E6E7 cells were arrested in G1 phase with 25 µM Lovastatin (Sigma) and released by exchanging the growth media with fresh media supplemented with 2.5 mM mevalonic acid (Sigma) as previously described (Javanmoghadam-Kamrani and Keyomarsi, 2008).

### Sample preparation for super-resolution microscopy

Samples were prepared for super-resolution imaging of telomere macromolecular structure as described elsewhere (Doksani et al., 2013). Cross-linking in this method is a modification of protocols used previously to visualize t-loops by electron microscopy (Cesare and Griffith, 2004; Griffith et al., 1999). Briefly, 1×10^7^ cells were washed in ice-cold 1x PBS and re-suspended in ice-cold lysis buffer (12.5 mM Tris pH 7.4, 5 mM KCl, 0.1 mM spermine, 0.25 mM spermidine, 175 mM sucrose) supplemented with protease inhibitor cocktail (Roche). The material was incubated for 10 minutes on ice, followed by addition of 0.03% (v/v) 10% NP-40 (Sigma) and incubation on ice for an additional 5 minutes. Cells were pelleted at 1000 g for 5 minutes at 4°C and washed once with ice-cold nuclei wash buffer (10 mM Tris-HCl pH 7.4, 15 mM NaCl, 60 mM KCl, 5 mM EDTA, 300 mM sucrose). The material was re-suspended in 3 ml of nuclei wash buffer in a 6-well non-tissue culture treated plate and incubated for 5 min while stirring on ice, in the dark, in the presence of 100 µg/ml Trioxsalen (Sigma) diluted in DMSO. The material was then exposed to 365 nm UV light, 2-3 cm from the light source (model UVL-56, UVP) for 30 min, while incubated on ice with continuous stirring. After crosslinking, the material was centrifuged at 1000 g for 5 min, washed with ice-cold nuclei wash buffer, and resuspended at 1×10^7^ cells/ml. The material was diluted 1:10 in spreading buffer (10 mM Tris-HCl 7.4, 10 mM EDTA, 0.05% SDS, 1 M NaCl) pre-warmed to 37°C and 100 µL of the suspension was immediately deposited on a 18 × 18 mm 170 µm thick coverslip using a Cellspin1 (Tharmac) at 600 rpm for 1 minute. Coverslips were fixed in −20°C methanol for 10 minutes then for 1 minute in −20°C acetone. Coverslips were briefly washed in room temperature 1x PBS then dehydrated through a 70%, 95% and 100% ethanol series, by incubating for 3 min in each condition.

### Telomere fluorescent *in situ* hybridization (FISH) for super-resolution microscopy

Ethanol dehydrated coverslips were denatured for 10 minutes at 80°C in the presence of C-rich telomere peptide nucleic acid (PNA) probe conjugated to Alexa fluor 647 (Alexa647-OO-ccctaaccctaaccctaa, Panagene) for STORM and STED, or Alexa fluor 488 (Alexa488-OO-ccctaaccctaaccctaa, Panagene) for SIM and Airyscan. PNA probe was prepared and diluted to 0.3 ng/ml as described previously (Cesare et al., 2015). Following hybridization overnight in a dark humidified box, coverslips were washed twice for 10 min in PNA Wash A (70% Formamide; 10 mM Tris-HCL pH 7.5) and thrice for five minutes in PNA Wash B (50 mM Tris-HCL pH 7.5, 150 mM NaCl, 0.8% Tween-20), all with gentle shaking. Coverslips were rinsed in MilliQ water and dehydrated through a 70%, 95%, and 100% ethanol series. Coverslips were air dried and stored dehydrated until imaging.

### STORM imaging

Coverslips were rehydrated in 1x PBS and stained with YoYo-1 at 0.1 µM for 10 minutes then washed twice for 10 minutes in 1x PBS. Coverslips were mounted air-sealed in a 1-well 18 × 18 mm Chamlide in the presence of an oxygen-scavenging 1x PBS solution supplemented with 25 mM HEPES, 25 mM glucose, 5% glycerol, 0.05 mg/ml glucose oxidase, 0.025 mg/ml horseradish peroxidase, 75 mM cysteamine. Total internal reflection fluorescence (TIRF) imaging of YoYo-1 labeled chromatin was used to identify regions where chromatin was deposited onto cover slips. STORM images were acquired under TIRF illumination on an ELYRA microscope (ZEISS) with a 1.46 NA alpha Plan-Apochromat 100x oil immersion objective. 15-30 mW from a 642 nm laser was used at the start of image capture. Continuous photo switching events per frame were collected and maintained by increasing the laser strength during image acquisition to counter balance photo-bleaching. 15,000-20,000 images were acquired per region with an exposure time of 35 ms using a cooled electron-multiplying CCD camera (Andor iXon DU-897D). Sample drift during image acquisition was corrected relative surface-immobilized 100 nm colloidal gold beads (BBInternational) placed on each sample. Images were captured in 512 × 512 pixels at 51.1 × 51.1 µm to scale and processed using ZEN software (ZEISS).

### SIM imaging

Slides were mounted in Prolong Gold (Life Technologies) in the presence of DAPI. Three dimensional-structured illumination microscopy (3D-SIM) was performed with a V3 DeltaVision OMX 3D-SIM fitted with a Blaze Module (GE Healthcare) and an Olympus UPlanSApo N 60x 1.42 NA oil objective (Olympus). Chromatin regions were identified by DAPI staining (405 nm laser) and telomere PNA images were captured on Edge scientific CMOS 512 × 512 pixel cameras (pco, Kelheim). Unprocessed image stacks were composed of 15 images per z-section(Strauss et al., 2012). The z-sections were completed at a spacing of every 125 nm for a total raw data image count of 120 images per 1 µm sample z-stack. The output of raw data from the OMX Blaze were then reconstructed to extract finer detail using the Gustafsson algorithms (Gustafsson et al., 2008) and SoftWorX software (GE Healthcare). Reconstructed images were rendered in 3D, with interpolation, using Imaris version 8.0.0 (Bitplane).

### STED imaging

Slides were mounted in Prolong Diamond (Life Technologies) in the presence of 0.1 µM YoYo-1. STED imaging was performed on a Leica SP8 gated STED 3X microscope with internal Leica HyD hybrid detector fitted with a HC PL APO 100x NA 1.40 Oil CS2 oil objective. Areas of chromatin labelled with YoYo-1 counter stain were localized using conventional resolution microscopy. Alexa Fluor 647 labelled telomeres were excited with 10-30% excitation power of a pulsed white light laser (1.5 mW) at 647nm wavelength and depleted with a continuous wave 775 nm STED laser (1.5W) with a typical maximum power of 30-60% at the back aperture of the objective. Images were captured in 1024 × 1024 pixels at 46.48 µm x 46.48 µm to scale, with a line averaging and frame accumulation setting of 6, and line accumulation and frame averaging of 1. STED images were exported using Leica Las X software and deconvolved using Huygens Professional software on default deconvolution settings based on the metadata.

### Airyscan imaging

Slides were mounted in Prolong Gold (Life Technologies) in the presence of DAPI. Airyscan imaging was performed on a ZEISS LSM880 AxioObserver confocal fluorescent microscope fitted with an Airyscan detector and a Plan-Apochromat 63x 1.4 NA M27 oil objective. Chromatin was identified with DAPI using conventional resolution microscopy prior to Airyscan imaging. Alexa Fluor 488 labelled telomeres were captured with 4% excitation power of 488 nm laser, 1X1 binning, detector gain of 950 and digital gain of 1 in super-resolution mode. A total of 5 z-stacks (200 nm) were captured with frame scanning mode, unidirectional scanning and line averaging of 2 in 1024 × 1024 pixels at 89.88 µm x 89.88 µm to scale. Z-stacks were Airyscan processed using batch mode in Zen Black software (ZEISS).

### Super-resolution microscopy scoring criteria

Images obtained from super resolution microscopy after processing were exported to Image J version 1.51H as TIF images with maintained scales. Each image was manually quantified with researchers blinded to experimental conditions, and the telomeres scored as looped, linear or ambiguous. We classified molecules as t-loops when we could discern an individual molecule consisting of a closed loop structure with a single attached tail, containing no gaps of PNA staining in that molecule of > 1 µm. A molecule was classified as linear when it was an individual molecule with two observable ends, containing no loops or branched structures, and no gaps in PNA staining of > 1 µm. Ambiguous molecules were all molecules that did not conform to the looped or linear definition. Areas of coverslips with densely packed and overlapping telomere molecules were not scored. Each loop and linear molecule were measured for contour length using the Image J trace function.

### Cross-linking efficiency determination

The trioxsalen fixation protocol for super-resolution microscopy was followed through the trioxsalen cross-linking step. Crosslinked cells were then pelleted by centrifugation at 1500 g for 5 minutes at 4°C. The DNA was digested with 100U/L micrococcal nuclease (NEB) at 37 °C for 10 min, and the reaction stopped with ice-cold nuclei wash buffer and centrifuged at 1500 g for 5 minutes. Pellets were resuspended in 70 mM Tris-HCl pH 8.0, 1 mM EDTA, 1.5% (v/v) SDS with 20 mg/mL proteinase K and incubated at 56°C for 1 hr. DNA was extracted with a Qiagen PCR purification kit according to manufacture instruction. Extracted DNA were separated on a 1% agarose gel and the di-nucleosome band isolated by gel-excision. DNA were purified using the Nucleospin Gel and PCR cleanup kit (Macherey-Nagel). Purified DNA were divided into two equal pools. One pool was heat denatured at 95°C for 5 min and snap cooled in ice-cold water. DNA samples were separated on a 1% agarose gel with ethidium bromide. Gel images were captured with an Alpha Innotech Fluorchem 5500 and band intensity quantified with Image J.

### Interphase-TIF, metaphase-TIF and cytogenetic assays

Interphase-TIF assays, metaphase-TIF assays, and cytogenetic chromosome preparations were performed as described elsewhere (Cesare et al., 2015). For metaphase-TIF assays, cells were centrifuged onto glass slides using a Tharmac Cellspin 1. In all assays, telomeres were visualized with FISH using an Alexa fluor 488 conjugated C-rich telomere PNA probe (Alexa488-OO-ccctaaccctaaccctaa, Panagene). For metaphase-TIF and cytogenetic chromosome spreads, images were captured using a ZEISS AxioImager Z.2 with a 63x, 1.4 NA oil objective and appropriate filter cubes, using a CoolCube1 camera (Metaystems). Automated metaphase finding and image acquisition for these experiments were done using the MetaSystems imaging platform as described elsewhere (Cesare et al., 2013). For interphase-TIF assays, images were captured using Zen software and a ZEISS AxioImager Z.2, with a 63x, 1.4 NA oil objective, appropriate filter cubes and an Axiocam 506 monochromatic camera (ZEISS).

### Live cell imaging

Mitotic duration and outcome were visualized with differential interference contrast microscopy on a ZEISS Cell Observer inverted wide field microscope, with 20x 0.8 NA air objective, at 37° C, 10% CO2 and atmospheric oxygen. Images were captured every six minutes, for a duration of 120 hours with an Axiocam 506 monochromatic camera and Zen software. Movies were compiled using Zen software and mitotic duration scored by eye from nuclear envelope breakdown until cytokinesis.

### Flow cytometry

Cells were collected with trypsin and the reaction quenched with growth media containing serum, then centrifuged and washed in 1x PBS. To determine genome content, cells were suspended in 1 ml of 1x PBS and fixed by dropwise addition of 3 ml of 100% ethanol while gently vortexing. Cells were fixed overnight at 4°C, washed twice in 1x PBS + 1% (w/v) BSA, then resuspended in 1x PBS supplemented with 1% BSA (w/v), 3.8 mM Tri-Sodium Citrate, 40 µg/ml Propidium Iodide and 50 μg/ml RNaseA. Samples were incubated at room temp for 30 min, then stored overnight at 4°C and analyzed the following day. For determination of H3-ser10 positive cells or γ-H2AX signal intensity, cells were collected with trypsin and washed in 1x PBS as described above, then aliquoted in batches of 1×106 cells and collected by centrifugation. Cell pellets were resuspended in 1 ml of 1x PBS + 2% formaldehyde (v/v) and fixed at 37°C for 10 min, chilled on ice for one minute, then permeablized with 9 ml of ice-cold 100% methanol by adding dropwise while gently vortexing. Samples were stored overnight at 4° C. Cells were washed twice in 1x PBS+ 1% (w/v) BSA, and blocked in this same solution for 10 min at RT, then pelleted with centrifugation. Cells were resuspended in 100 µl 1x PBS + 1% BSA (w/v) supplemented with 20 µg/mL primary antibody (either unconjugated anti-γ-H2AX, or Alexa488 conjugated anti-H3ser10) and incubated for 1 hour at RT with shaking. Cells were washed thrice for 5 minutes in PBS with 1% BSA. For γ-H2AX, the process of antibody staining and washing was repeated with a secondary Alexa Fluor 488 conjugated goat anti-mouse antibody at 40 µg/mL for 1 hour. After final wash, cells were stained with propidium iodide and RNaseA treated as described above. Flow cytometry was performed on a FACSCanto with FACSDiva software V6.1.3 (Becton Dickson) FSC and SSC were used to gate for the cell population (P1) and single cells were identified by excluding cell doublets through comparison of propidium iodide area and height (P2). At least 10,000 cells/replicates were collected in the P2 gate for all experiments. Data were exported to FlowJo Version 10 for analysis. For γ-H2AX staining experiments, the G1 population was gated as identified in Supplementary Figure 8. For H3-ser10 staining experiments, mitotic cells were identified as described previously(Hayashi et al., 2012).

### Western Blots

Cells were collected with trypsin and the reaction quenched with growth media containing serum. Cells were washed in 1x PBS and homogenized in NuPage 4x LDS sample buffer without EDTA (Invitrogen) supplemented with 2% (v/v) β-mercaptoethanol (Sigma) and 5% (v/v) Benzonase (Merck Millipore) at 1×104 cells/µL for 1 hr at room temperature. Cell lysate was denatured at 68°C for 10 minutes prior to resolution at 10 µL/well on Nu-PAGE 4-12% Bis-Tris gradient gels according to manufacturer’s protocols. Protein was transferred to nitrocellulose (Amersham) at 100 V for 1 hr and the membranes blocked for 1 hr with 5% (w/v) powdered skim milk in 1x PBS + 0.1% (v/v) Tween 20 (1x PBST). Membranes were probed overnight with primary antibody diluted in 1x PBST + 5% (w/v) powdered skim milk at 4°C with gentle nutation. The following morning blots were washed three times for 10 minutes with 1x PBST then probed with Horse Radish Peroxidase-conjugated secondary antibody diluted in 1x PBST + 5% (w/v) powdered skim milk for one hour at room temperature, followed by three washes in 1x PBST. Membranes were rinsed in deionized water blotted dry and overlaid with 0.22 µm filtered Enhanced Chemiluminescent western blotting substrate for 5 minutes (Pierce) before visualization with a Fujifilm LAS 4000 luminescent image analyzer and associated software (GE Health).

### Telomere restriction fragment analysis

Telomere restriction fragments (TRFs) were prepared by *Hinf*I and *Rsa*I digestion of genomic DNA, and 2 µg of digested DNA separated by one dimensional pulsed-field gel electrophoresis, or 20 µg by two-dimensional pulsed field (1st dimension) and standard (2^nd^ dimension) gel electrophoresis as described previously (Pickett et al., 2011). Native and denatured telomere signal was then probed with a [γ-^32^P]-ATP-labelled (CCCTAA)^3^ oligonucleotide probe and detected by PhosphorImager screens as described elsewhere (Pickett et al., 2011). PhosphorImager screens were imaged on a Typhoon FLA 9500 and quantitated on ImageQuantTL software (GE Healthcare Life Sciences). Total single strand G-rich telomere signal was measured per lane in all samples from ∼ 2 to 150 kb using an identical sized ROI. The amount of single strand G-rich telomere signal was quantified relative to total telomere signal in the denatured gel and normalized to the 0 hour 4-OHT sample.

### Antibodies

Primary antibodies used in this study: Actin (Sigma, A2228), CHK2 (Millipore, 05-649), Alexa Fluor 488 conjugated H3-ser10 (Cell Signaling Technology, 9708), γ-H2AX (Millipore 05-636), Myc-Tag (Cell Signaling Technology, 2276), TRF2 (Novus NB110-57130), and 53BP1 (Santa Cruz, sc-22760). We used highly cross-absorbed secondary antibodies conjugated to Alexa Fluor 488, Alexa Fluor 568 or Alexa Fluor 647 (Life Technologies) for immunofluorescence and secondary antibodies conjugated to Horse Radish Peroxidase (Dako) for Western blots.

### Statistics and data presentation

Statistical analysis was performed using GraphPad Prism Version 5. All box plots are displayed using the Tukey method where the box extends from the 25^th^ to the 75^th^ percentile data points and the line represents the mean. The upper whisker represents data points ranging up to the 75^th^ percentile + (1.5 × the inner quartile range), or the largest value data point if no data points are outside this range. The lower whisker represents data points ranging down to the 25^th^ percentile – (1.5 × the inner quartile range), or the smallest data point if no data points are outside this range. Data points outside these ranges are shown as individual points. Figures were prepared using Adobe Photoshop and Illustrator.

## SUPPLEMENTAL INFORMATION

### Telomere loops regulate ATM activation at chromosome ends

**Figure S1 related to Figure 1.**
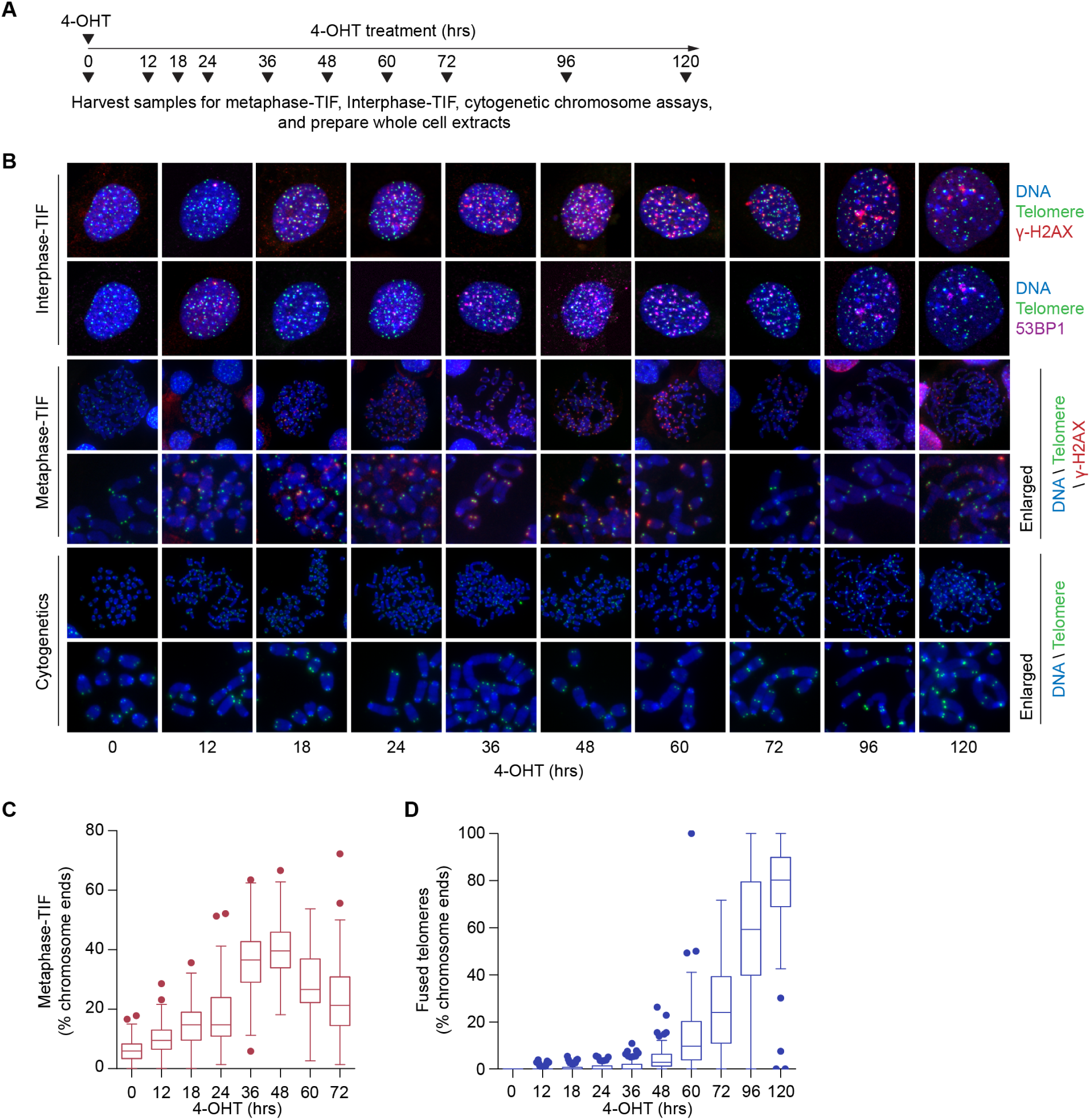
Mouse telomeres transition through a DDR-positive/NHEJ-resistant state following *TRF2* deletion. A) Timeline of the experimentation in Figure 1 and (B-D)below. B) Interphase-TIF, metaphase-TIF and cytogenetic chromosome preparations from *TRF2^F/F^ CreER LgT* cells after 4-OHT treatment. C, D) Quantitation of metaphase-TIF (C) and telomere fusions (D) in *TRF2^F/F^ CreER LgT* cells following *TRF2* deletion. These are a different representation of the same data from Figure 1B (all data points from n = 3 biological replicates of ≥ 30 mitotic spreads per replicate presented in a Tukey box plot).

**Figure S2 related to Figure 1.**
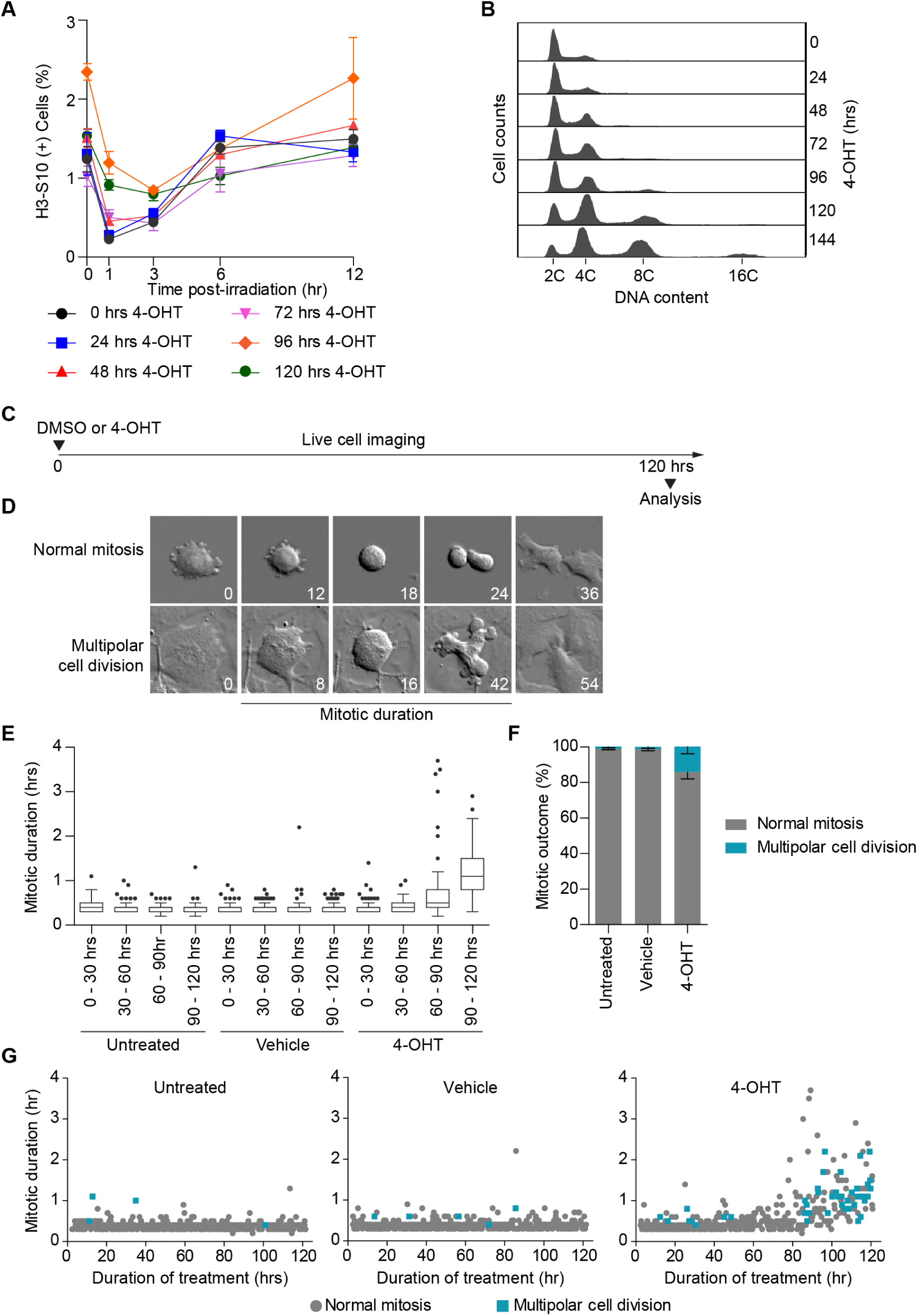
Phenotypic response to telomere deprotection in *TRF2F/F CreER LgT* MEFs. A) Mitotic content of *TRF2^F/F^ CreER LgT* cultures as determined by H3-S10 staining following TRF2 deletion, ± gamma-irradiation (IR). At the indicated time following 4-OHT treatment, *TRF2^F/F^ CreER LgT* MEFs were treated with 1 Gy IR and allowed to recover for 0, 1, 3, 6 or 12 hours. Mitotic cells were quantified by flow cytometry (mean ± s.e.m., n = 3 biological replicates). B) Genome content distribution as determined by flow cytometry in *TRF2^F/F^ CreER LgT* MEFs following *TRF2* deletion. C) Timeline of the live cell imaging experimentation in (D-G). D) Representative images of *TRF2^F/F^ CreER LgT* MEFs demonstrating mitotic duration and outcome as measured by live cell imaging. Time is shown in min relative to the first image.E)Quantitation of mitotic duration in untreated, vehicle (DMSO) and 4-OHT treated *TRF2^F/F^ CreER LgT* MEFs. Live cell imaging was carried out for 120 hours. Each point represents an individual mitosis. Mitotic events are grouped into 30 hour intervals (Pooled data from n = 3 biological replicates compiled into Tukey Box plots, ≥ 30 cell divisions per interval per replicate). Mitotic outcome of the cell divisions shown in (E) (mean ± s.e.m., n = 3 biological replicates of ≥ 120 cell divisions per replicate). G) All mitotic events from (E). Each point represents a mitotic event. Location of a symbol relative to the x-axis indicates the time after 4-OHT addition when that mitosis started. Height on the y-axis represents the mitotic duration of that mitosis. The symbol identifies mitotic outcome (pooled data from n = 3 biological replicates of ≥ 120 cell divisions per replicate).

**Figure S3 related to Figure 2.**
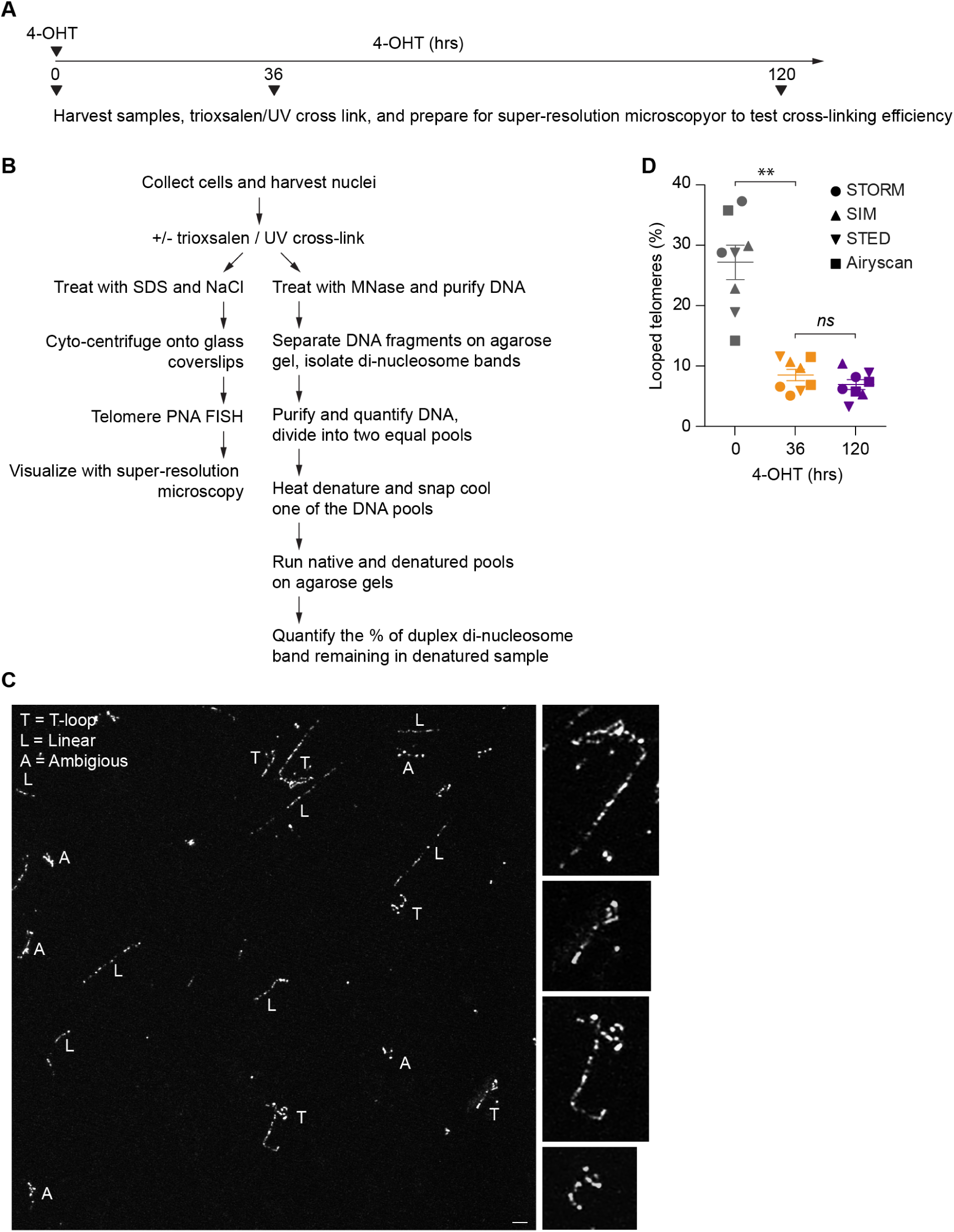
Experimental timeline, protocol, and scoring of telomere structure from *TRF2^F/F^ CreER LgT* MEFs using super-resolution microscopy. A) Experimental timeline of the experimentation in Figure 2A-E and below. B) Experimental protocol for preparing samples for super-resolution imaging of telomere macromolecular structure and determination of cross-linking efficiency (Doksani et al., 2013). This protocol was used for all super-resolution imaging and cross-linking efficiency experiments described in this study. C) Representative field and scoring of an Airyscan microscopy image of *TRF2^F/F^ CreER LgT* telomeres (bar = 1 µm). Expanded t-loops are shown on the right. Some of the expanded molecules have been rotated relative to the image on the left. D) Percentage of looped telomeres observed by super-resolution microscopy at the indicated times of 4-OHT. Calculations include ambiguous molecules in the data set (mean ± s.e.m., n = 8 biological replicates of ≥ 580 molecules per replicate, ** *p* < *0.01, ns* = not significant, student t-test).

**Figure S4 related to figure 2.**
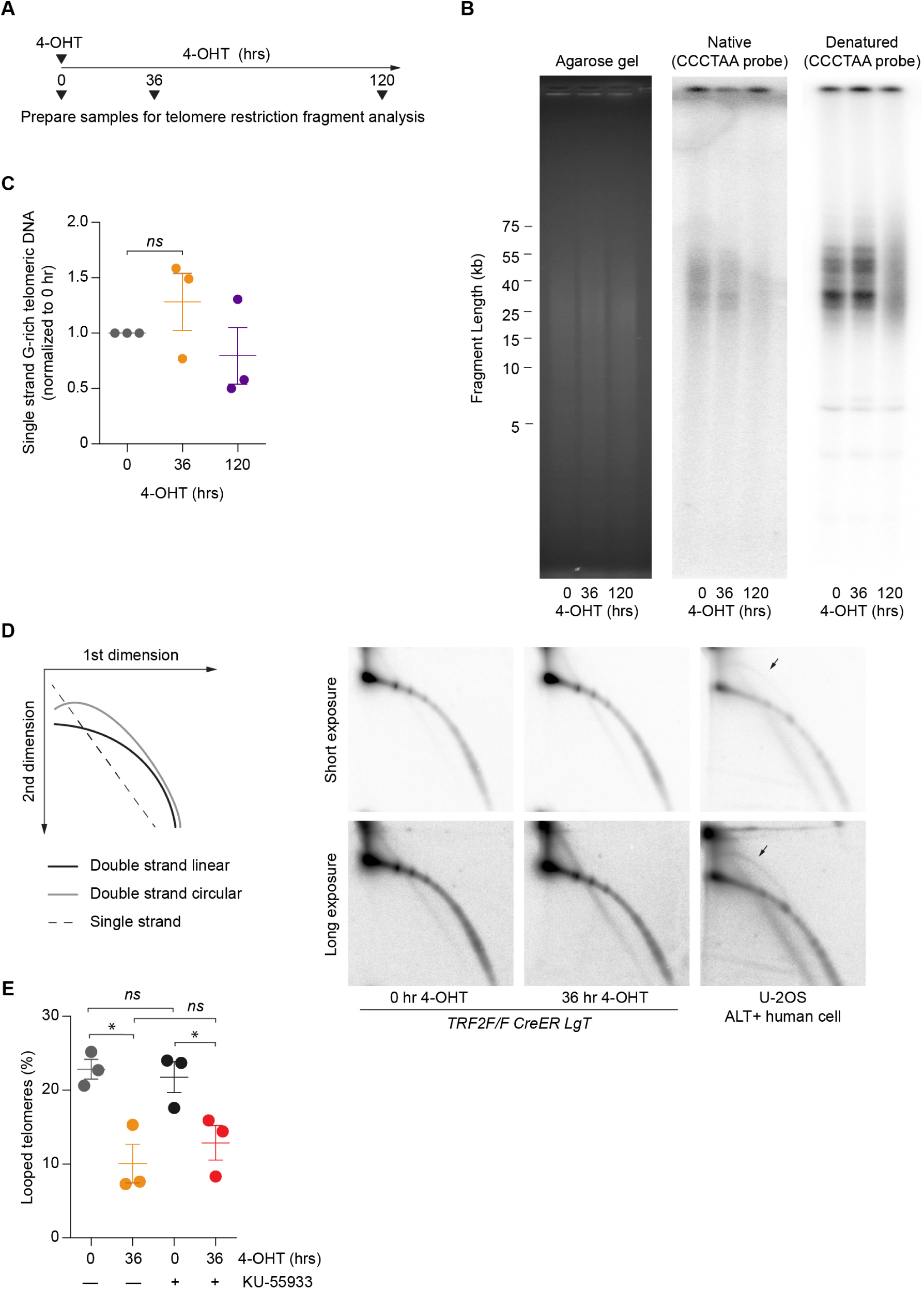
Telomere linearization results from t-loop unfolding independent of ATM activity. **A**) Experimental timeline of the experimentation below. B) Telomere restriction fragments were generated from *TRF2^F/F^ CreER LgT* MEFs after the indicated duration of 4-OHT treatment and separated by pulsed-field gel electrophoresis. The gel was stained with ethidium bromide (left), subjected to hybridization with a C-rich telomere probe under native conditions to detect single-strand G-rich telomeric DNA as a measure of the telomere overhang (middle), then denatured and re-hybridized with the same C-rich probe to identify double-strand telomeric DNA (right). C) Quantitation of the single-strand G-rich telomeric DNA signal from (B) (mean ± s.e.m., n = 3 biological replicates, *ns* = not significant, student t-test). D) Telomere restriction fragments from *TRF2^F/F^ CreER LgT* MEFs after the indicated duration of 4-OHT treatment were separated on 2D agarose gels and probed after denaturation with a C-rich telomere probe. U-2OS ALT-positive human cells are shown as a t-circle positive control (Cesare and Griffith, 2004). A diagram depicting the telomeric DNA arcs in the 2D gels is shown on the right. The t-circle arc in U-2OS cells is indicated by the arrow. E) Percentage of looped telomeres observed by Airyscan microscopy at the indicated times of 4-OHT treatment +/-KU-55933. Calculations include ambiguous molecules in the data set (mean ± s.e.m., n = 3 biological replicates of ≥ 630 molecules scored per replicate, * *p < 0.05, ns* = not significant, student t-test).

**Figure S5 related to figure 4.**
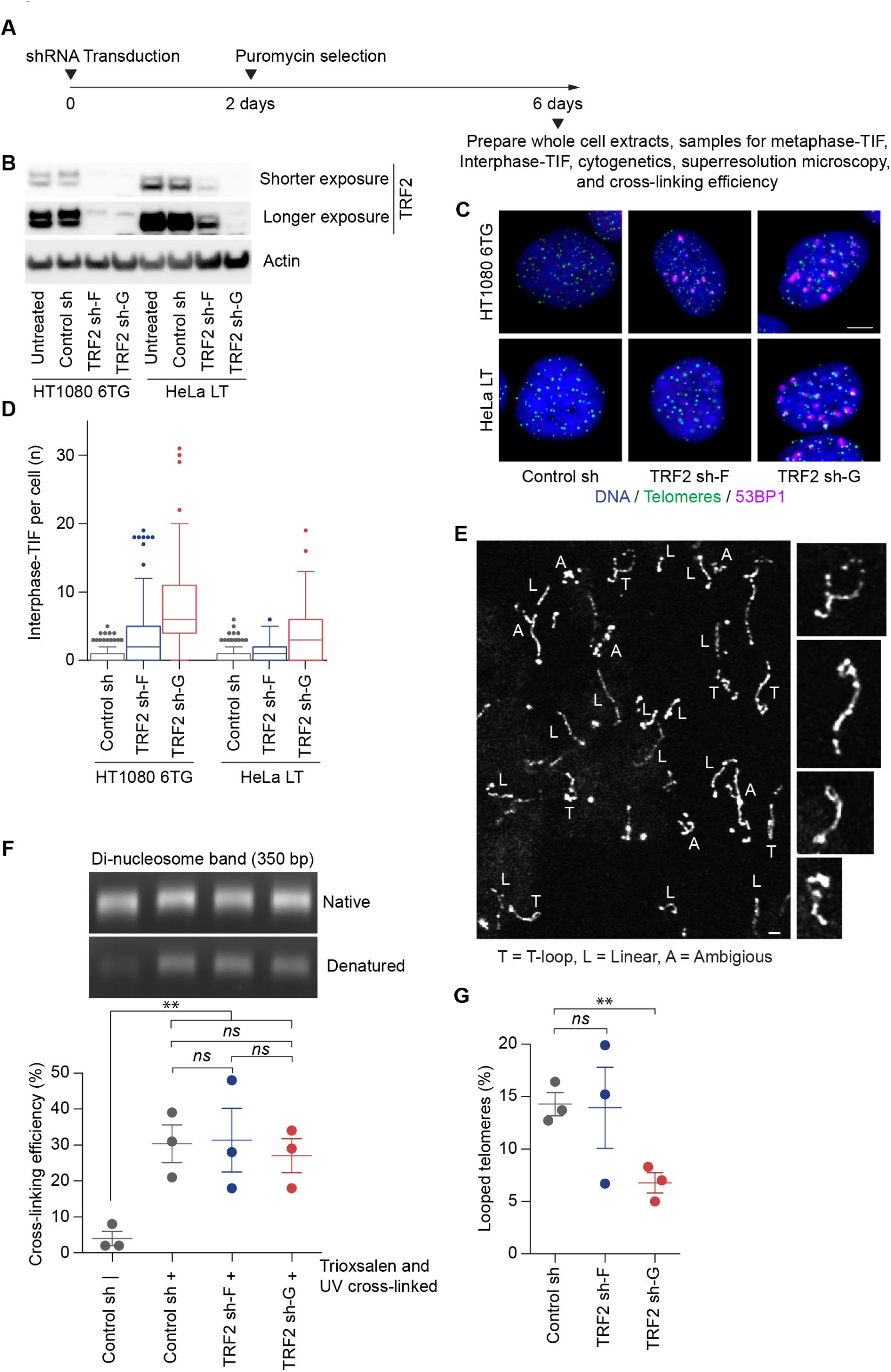
TRF2 depletion, cross-linking efficiency, and t-loop scoring in HeLa LT cells. A) Timeline of the experimentation in Figure 4B-F and (B-G) below. B) Western blots of whole cell extracts from HeLa LT and HT1080 6TG cells stably transduced with control and TRF2 shRNAs. C) Representative interphase-TIF images from HeLa LT and HT1080 6TG cells stably transduced with control and TRF2 shRNAs. D) Quantitation of interphase-TIF in HeLa LT and HT1080 6TG cells stably transduced with control and TRF2 shRNAs (n = 3 biological replicates scoring ≥ 50 nuclei per replicate are compiled into a Tukey box plot). E) Representative field and scoring of HeLa LT telomeres visualized by Airyscan microscopy (bar = 1 µm). Expanded t-loops are shown to the right. Some examples are rotated relative to the image on the left. F) Determination of cross-linking efficiency in HeLa LT cells stably transduced with control and TRF2 shRNAs (mean ± s.e.m., n = 3 biological replicates, ** *p < 0.01, ns* = not significant, student t-test). G) Quantitation of looped telomeres from HeLa LT observed by Airyscan microscopy. Calculations include ambiguous molecules from the data set (mean ± s.e.m., n = 3 biological replicates scoring ≥ 929 molecules per replicate, ** *p < 0.01*, ns = not significant, student t-test).

**Figure S6 related to figure 5.**
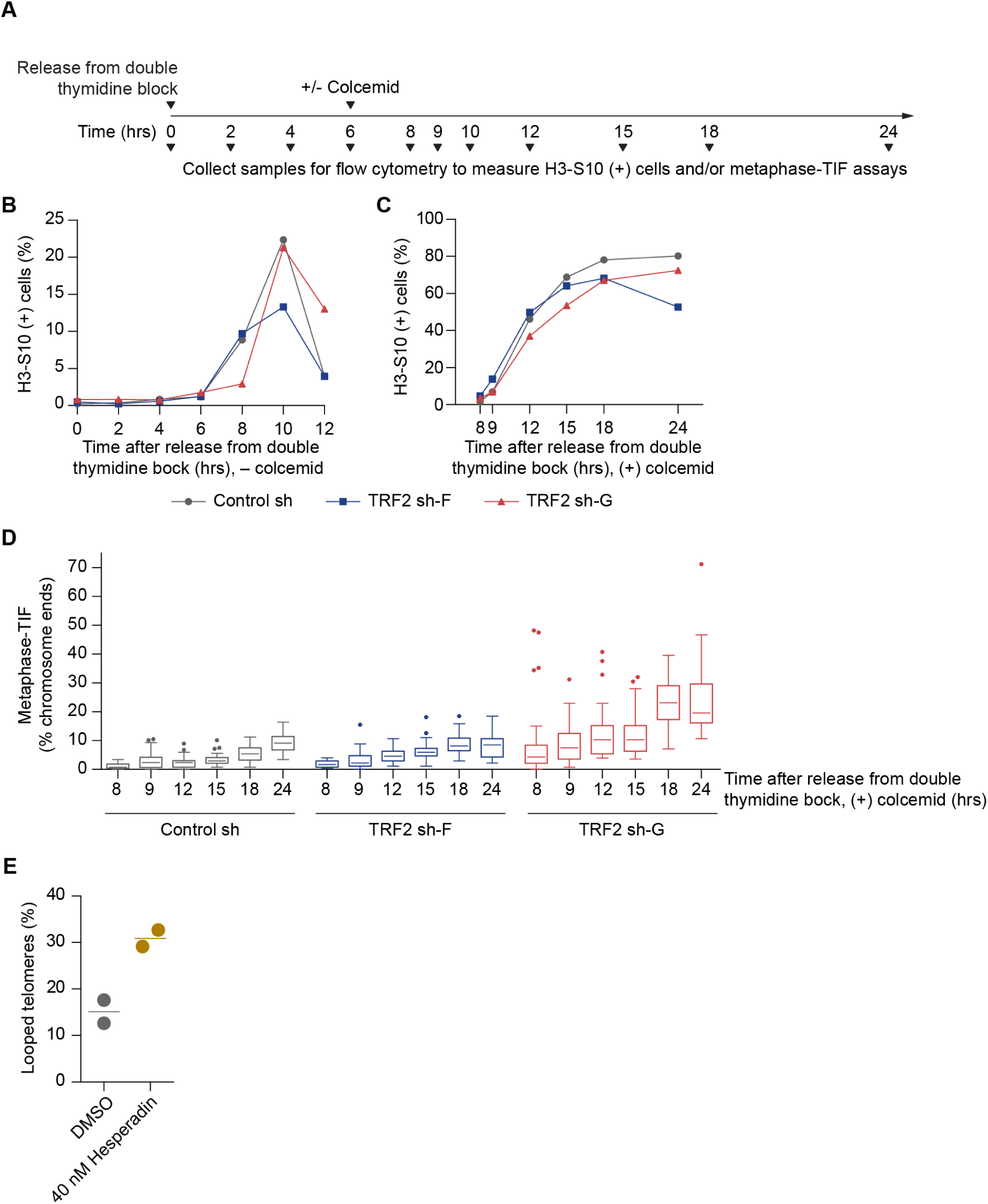
Experimental conditions to induce maximal mitotic telomere deprotection in HeLa LT cells. A) Timeline of the experimentation in (B-D) below. B) The percentage of mitotic cells in HeLa LT shRNA transduced cultures as determined by flow cytometry of H3-S10 positive cells following release from synchronization with a double thymidine bock. C) The percentage of mitotic cells as determined by flow cytometry of H3-S10 positive cells in HeLa LT shRNA transduced cultures following release from a double thymidine block and colcemid addition six hours later. D) Quantitation of metaphase-TIF assays in shRNA transduced HeLa LT cells for the conditions and time points shown in (C) (n ≥ 30 mitotic cells per condition. shown in a Tukey box plot). E) Quantitation of looped telomeres in mitotically arrested HeLa LT cells. Calculations include ambiguous molecules (n = 2 biological replicates of ≥ 956 molecules per replicate, line depicts the mean).

**Figure S7 related to Figures 6 and 7.**
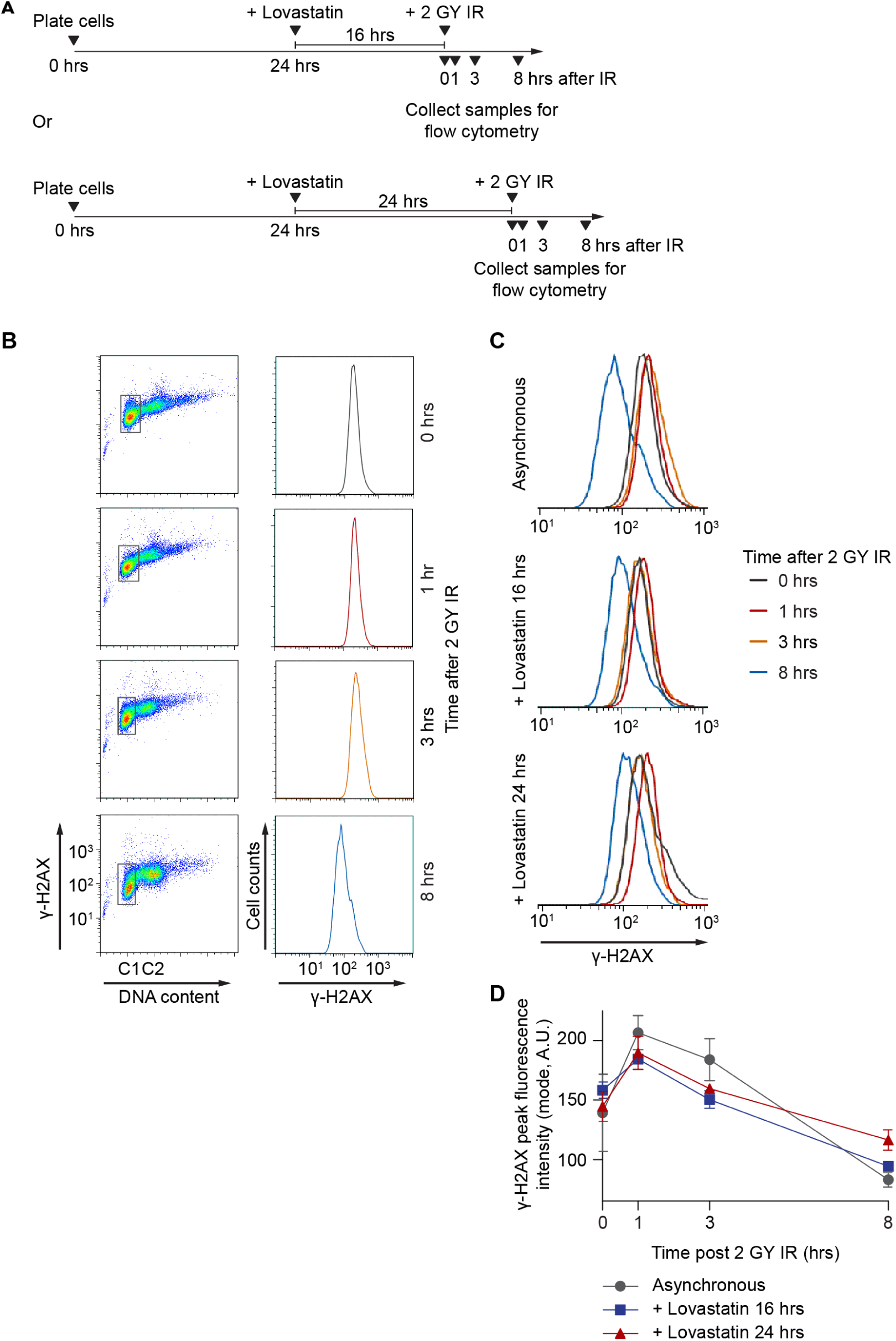
Double-strand break repair is active during Lovastatin induced G1-arrest. A) Timeline of the experimentation in (B-D) below. B) Example of γ-H2AX signal measurement by flow cytometry. At the indicated times following irradiation, cells were collected and stained for γ-H2AX, and with propidium iodide to identify DNA content. Flow cytometry data was gated on G1-phase cells (left) and γ-H2AX signal distribution determined for the gated cell population (right). C) Examples of γ-H2AX signal intensity in asynchronous and Lovastatin synchronized cells following IR and recovery. D) Quantitation of the experiment in (C) (mode + s.d., n = 3 biological replicates).

**Table S1:** Quantitative data from super-resolution imaging of murine telomeres

| Microscope | Tamoxifen (hr) | Additional treatment | Repeat | (n) |  |  |  |  | % including ambiguous |  |  | % excluding ambiguous |  |
| --- | --- | --- | --- | --- | --- | --- | --- | --- | --- | --- | --- | --- | --- |
|  |  |  |  | Total | Linear | Looped | Ambiguous | Total - Ambiguous | Linear | Looped | Ambiguous | Linear | Looped |
| STORM | 0 | none | 1 | 1600 | 401 | 597 | 602 | 998 | 25.1 | 37.3 | 37.6 | 40.2 | 59.8 |
| STORM | 0 | none | 2 | 736 | 286 | 212 | 238 | 498 | 38.9 | 28.8 | 32.3 | 57.4 | 42.6 |
| STED | 0 | none | 1 | 1517 | 449 | 437 | 631 | 886 | 29.6 | 28.8 | 41.6 | 50.7 | 49.3 |
| STED | 0 | none | 2 | 2201 | 1121 | 417 | 663 | 1538 | 50.9 | 18.9 | 30.1 | 72.9 | 27.1 |
| SIM | 0 | none | 1 | 815 | 380 | 186 | 249 | 566 | 46.6 | 22.8 | 30.6 | 67.1 | 32.9 |
| SIM | 0 | none | 2 | 1239 | 283 | 371 | 585 | 654 | 22.8 | 29.9 | 47.2 | 43.3 | 56.7 |
| Airyscan | 0 | none | 1 | 4609 | 1327 | 1652 | 1630 | 2979 | 28.8 | 35.8 | 35.4 | 44.5 | 55.5 |
| Airyscan | 0 | none | 2 | 2538 | 728 | 361 | 1449 | 1089 | 28.7 | 14.2 | 57.1 | 66.9 | 33.1 |
| STORM | 36 | none | 1 | 1049 | 409 | 53 | 587 | 462 | 39.0 | 5.1 | 56.0 | 88.5 | 11.5 |
| STORM | 36 | none | 2 | 1289 | 645 | 85 | 559 | 730 | 50.0 | 6.6 | 43.4 | 88.4 | 11.6 |
| STED | 36 | none | 1 | 1685 | 633 | 196 | 856 | 829 | 37.6 | 11.6 | 50.8 | 76.4 | 23.6 |
| STED | 36 | none | 2 | 1154 | 753 | 68 | 333 | 821 | 65.3 | 5.9 | 28.9 | 91.7 | 8.3 |
| SIM | 36 | none | 1 | 1051 | 639 | 112 | 300 | 751 | 60.8 | 10.7 | 28.5 | 85.1 | 14.9 |
| SIM | 36 | none | 2 | 1403 | 484 | 136 | 783 | 620 | 34.5 | 9.7 | 55.8 | 78.1 | 21.9 |
| Airyscan | 36 | none | 1 | 3313 | 1450 | 382 | 1481 | 1832 | 43.8 | 11.5 | 44.7 | 79.1 | 20.9 |
| Airyscan | 36 | none | 2 | 3833 | 1465 | 263 | 2105 | 1728 | 38.2 | 6.9 | 54.9 | 84.8 | 15.2 |
| STORM | 120 | none | 1 | 919 | 771 | 75 | 73 | 846 | 83.9 | 8.2 | 7.9 | 91.1 | 8.9 |
| STORM | 120 | none | 2 | 1068 | 905 | 66 | 97 | 971 | 84.7 | 6.2 | 9.1 | 93.2 | 6.8 |
| STED | 120 | none | 1 | 1094 | 528 | 97 | 469 | 625 | 48.3 | 8.9 | 42.9 | 84.5 | 15.5 |
| STED | 120 | none | 2 | 1364 | 1131 | 45 | 188 | 1176 | 82.9 | 3.3 | 13.8 | 96.2 | 3.8 |
| SIM | 120 | none | 1 | 580 | 418 | 31 | 131 | 449 | 72.1 | 5.3 | 22.6 | 93.1 | 6.9 |
| SIM | 120 | none | 2 | 1060 | 579 | 110 | 371 | 689 | 54.6 | 10.4 | 35.0 | 84.0 | 16.0 |
| Airyscan | 120 | none | 1 | 2343 | 1465 | 173 | 705 | 1638 | 62.5 | 7.4 | 30.1 | 89.4 | 10.6 |
| Airyscan | 120 | none | 2 | 1896 | 956 | 110 | 830 | 1066 | 50.4 | 5.8 | 43.8 | 89.7 | 10.3 |
| Airyscan | 0 | DMSO | 1 | 1536 | 827 | 316 | 393 | 1143 | 53.8 | 20.6 | 25.6 | 72.4 | 27.6 |
| Airyscan | 0 | DMSO | 2 | 824 | 503 | 187 | 134 | 690 | 61.0 | 22.7 | 16.3 | 72.9 | 27.1 |
| Airyscan | 0 | DMSO | 3 | 820 | 485 | 207 | 128 | 692 | 59.1 | 25.2 | 15.6 | 70.1 | 29.9 |
| Airyscan | 36 | DMSO | 1 | 1374 | 868 | 210 | 296 | 1078 | 63.2 | 15.3 | 21.5 | 80.5 | 19.5 |
| Airyscan | 36 | DMSO | 2 | 630 | 478 | 46 | 106 | 524 | 75.9 | 7.3 | 16.8 | 91.2 | 8.8 |
| Airyscan | 36 | DMSO | 3 | 1074 | 830 | 82 | 162 | 912 | 77.3 | 7.6 | 15.1 | 91.0 | 9.0 |
| Airyscan | 0 | 1.5 μM KU-55933 | 1 | 1639 | 965 | 288 | 386 | 1253 | 58.9 | 17.6 | 23.6 | 77.0 | 23.0 |
| Airyscan | 0 | 1.5 μM KU-55933 | 2 | 708 | 437 | 170 | 101 | 607 | 61.7 | 24.0 | 14.3 | 72.0 | 28.0 |
| Airyscan | 0 | 1.5 μM KU-55933 | 3 | 1004 | 647 | 238 | 119 | 885 | 64.4 | 23.7 | 11.9 | 73.1 | 26.9 |
| Airyscan | 36 | 1.5 μM KU-55933 | 1 | 1054 | 714 | 88 | 252 | 802 | 67.7 | 8.3 | 23.9 | 89.0 | 11.0 |
| Airyscan | 36 | 1.5 μM KU-55933 | 2 | 699 | 442 | 111 | 146 | 553 | 63.2 | 15.9 | 20.9 | 79.9 | 20.1 |
| Airyscan | 36 | 1.5 μM KU-55933 | 3 | 819 | 550 | 118 | 151 | 668 | 67.2 | 14.4 | 18.4 | 82.3 | 17.7 |
| Airyscan | 0 | nMyc-TRF2 | 1 | 1984 | 1475 | 364 | 145 | 1839 | 74.3 | 18.3 | 7.3 | 80.2 | 19.8 |
| Airyscan | 0 | nMyc-TRF2 | 2 | 1044 | 763 | 173 | 108 | 936 | 73.1 | 16.6 | 10.3 | 81.5 | 18.5 |
| Airyscan | 0 | nMyc-TRF2 | 3 | 1098 | 788 | 213 | 97 | 1001 | 71.8 | 19.4 | 8.8 | 78.7 | 21.3 |
| Airyscan | 120 | nMyc-TRF2 | 1 | 1696 | 1074 | 423 | 199 | 1497 | 63.3 | 24.9 | 11.7 | 71.7 | 28.3 |
| Airyscan | 120 | nMyc-TRF2 | 2 | 1809 | 1257 | 406 | 146 | 1663 | 69.5 | 22.4 | 8.1 | 75.6 | 24.4 |
| Airyscan | 120 | nMyc-TRF2 | 3 | 1768 | 1263 | 360 | 145 | 1623 | 71.4 | 20.4 | 8.2 | 77.8 | 22.2 |
| Airyscan | 0 | nMyc-TRF<sup>CT</sup> | 1 | 827 | 488 | 225 | 114 | 713 | 59.0 | 27.2 | 13.8 | 68.4 | 31.6 |
| Airyscan | 0 | nMyc-TRF<sup>CT</sup> | 2 | 911 | 649 | 133 | 129 | 782 | 71.2 | 14.6 | 14.2 | 83.0 | 17.0 |
| Airyscan | 0 | nMyc-TRF<sup>CT</sup> | 3 | 1072 | 751 | 153 | 168 | 904 | 70.1 | 14.3 | 15.7 | 83.1 | 16.9 |
| Airyscan | 120 | nMyc-TRF<sup>CT</sup> | 1 | 942 | 842 | 46 | 54 | 888 | 89.4 | 4.9 | 5.7 | 94.8 | 5.2 |
| Airyscan | 120 | nMyc-TRF<sup>CT</sup> | 2 | 1047 | 959 | 53 | 35 | 1012 | 91.6 | 5.1 | 3.3 | 94.8 | 5.2 |
| Airyscan | 120 | nMyc-TRF<sup>CT</sup> | 3 | 919 | 823 | 49 | 47 | 872 | 89.6 | 5.3 | 5.1 | 94.4 | 5.6 |

**Supplementary Table 2:**
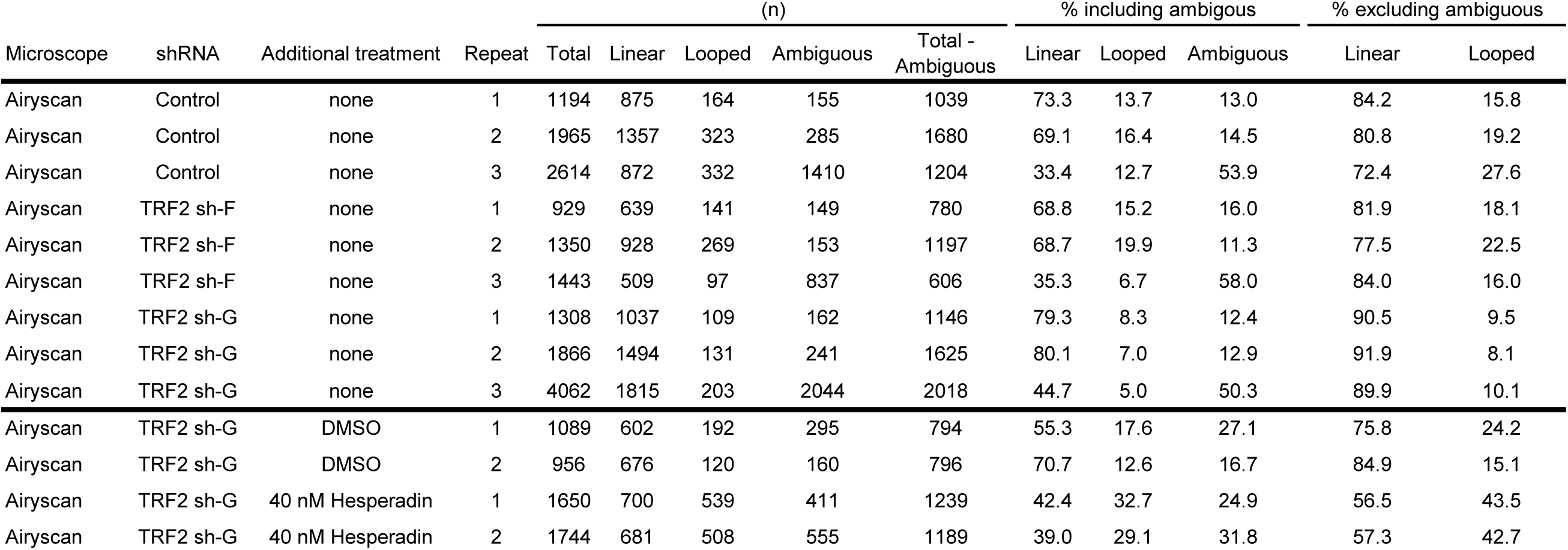
Quantitative data from super-resolution imaging of human telomeres

